# The MuSK-BMP pathway regulates synaptic Nav1.4 localization and muscle excitability

**DOI:** 10.1101/2023.10.24.563837

**Authors:** L. A. Fish, M. D. Ewing, D. Jaime, K. A. Rich, C. Xi, X. Wang, R. E. Feder, K. A. Wharton, M. M. Rich, W. D. Arnold, J. R. Fallon

**Author notes:** Denotes corresponding authors. Email addresses. **Conflict of interest:** LAF, DJ and JRF are co-inventors on patents to Brown University regarding the MuSK-BMP pathway. JRF is a co-founder of Bolden Therapeutics, to which these patents are licensed.

## Abstract

The neuromuscular junction (NMJ) is the linchpin of nerve-evoked muscle contraction. Broadly considered, the function of the NMJ is to transduce a nerve action potential into a muscle fiber action potential (MFAP). Efficient information transfer requires both cholinergic signaling, responsible for the generation of endplate potentials (EPPs), and excitation, the activation of postsynaptic voltage-gated sodium channels (Nav1.4) to trigger MFAPs. In contrast to the cholinergic apparatus, the signaling pathways that organize Nav1.4 and muscle fiber excitability are poorly characterized. Muscle-specific kinase (MuSK), in addition to its Ig1 domain-dependent role as an agrin-LRP4 receptor, is also a BMP co-receptor that binds BMPs via its Ig3 domain and shapes BMP-induced signaling and transcriptional output. Here we probed the function of the MuSK-BMP pathway at the NMJ using mice lacking the MuSK Ig3 domain (‘ΔIg3-MuSK’). Synapses formed normally in ΔIg3-MuSK animals, but the postsynaptic apparatus was fragmented from the first weeks of life. Anatomical denervation was not observed at any age examined. Moreover, spontaneous and nerve-evoked acetylcholine release, AChR density, and endplate currents were comparable to WT. However, trains of nerve-evoked MFAPs in ΔIg3-MuSK muscle were abnormal as revealed by increased jitter and blocking in single fiber electromyography. Further, nerve-evoked compound muscle action potentials (CMAPs), as well as twitch and tetanic muscle torque force production, were also diminished. Finally, Nav1.4 levels were reduced at ΔIg3-MuSK synapses but not at the extrajunctional sarcolemma, indicating that the observed excitability defects are the result of impaired localization of this voltage-gated ion channel at the NMJ. We propose that MuSK plays two distinct roles at the NMJ: as an agrin-LRP4 receptor necessary for establishing and maintaining cholinergic signaling, and as a BMP co-receptor required for maintaining proper Nav1.4 density, nerve-evoked muscle excitability and force production. The MuSK-BMP pathway thus emerges as a target for modulating excitability and functional innervation, which are defective in conditions such as congenital myasthenic syndromes and aging.

**Significance Statement:** The neuromuscular junction (NMJ) is required for nerve-evoked muscle contraction and movement, and its function is compromised during aging and disease. Although the mechanisms underlying neurotransmitter release and cholinergic response at this synapse have been studied extensively, the machinery necessary for nerve-evoked muscle excitation are incompletely characterized. We show that MuSK (Muscle-specific kinase), in its role as a BMP co-receptor, regulates NMJ structure as well as the localization of the voltage-gated sodium channels necessary for full nerve-evoked muscle fiber excitation and force production. This novel function of MuSK is structurally and mechanistically distinct from its role in organizing cholinergic machinery. The MuSK-BMP pathway thus presents a new opportunity to understand mechanisms that may preserve or enhance neuromuscular excitability in the face of aging and disease.

## Introduction

The neuromuscular junction (NMJ) is the highly specialized synapse between motor neuron and myofiber that transduces a nerve action potential into a muscle fiber action potential (MFAP), thus enabling nerve-evoked muscle contraction (Fatt and Katz, 1952; Sanes and Lichtman, 1999; Slater, 2017). The postsynaptic membrane harbors high densities (∼10,000/µ^2^) of acetylcholine receptors (AChRs) that generate endplate potentials (EPPs) in response to acetylcholine (ACh) released by the nerve terminal. EPPs are in turn amplified by subadjacent voltage-gated Nav1.4 sodium channels, also present at high density in the postsynaptic membrane, to generate MFAPs (Sanes and Lichtman, 2001; Slater, 2008). MFAP generation depends on excitability of the postsynaptic region, mediated by the proper localization and function of Nav1.4 channels (Slater, 2008; Schiaffino and Reggiani, 2011).

NMJs are remarkably stable, with the AChR-rich postsynaptic domains largely continuous and changing little over much of life (Li et al., 2011). However, these domains become fragmented in diseases such as muscular dystrophy and ALS as well as during aging (Valdez et al., 2012; Haddix et al., 2018; Belhasan and Akaaboune, 2020; Fish and Fallon, 2020). In some of these settings, this fragmentation is accompanied by nerve terminal loss (‘anatomical denervation’); however, in many settings pre- and post-synaptic apposition is sustained at fragmented synapses and cholinergic signaling is largely normal (Valdez et al., 2010; Poort et al., 2016; Willadt et al., 2016; Slater, 2020). Notably, in aged muscle, nerve-evoked excitability and muscle contraction can be compromised even when cholinergic signaling is unaffected (Chugh et al., 2020). These observations suggest that distinct molecular mechanisms may regulate innervation, cholinergic function and excitability.

Muscle-specific kinase (MuSK) plays a central role in configuring the NMJ. MuSK is a receptor tyrosine kinase containing three immunoglobulin-like (Ig) and a Crd/Fz domain extracellularly (Fish and Fallon, 2020). The best-known function of MuSK is in the formation and maintenance of the apparatus supporting cholinergic signaling. In this context MuSK engages agrin-LRP4 via its Ig1 domain with consequent activation of its tyrosine kinase. This pathway is necessary for the formation and maintenance of AChR-rich postsynaptic domains and the proper positioning of the nerve terminal in development (Glass et al., 1996; Watty et al., 2000; Stiegler et al., 2006; Zhang et al., 2011; Zong et al., 2012; Huijbers et al., 2013). However, whether MuSK plays roles in the assembly of other NMJ components has not been determined.

We recently reported that MuSK is also a bone morphogenetic protein (BMP) co-receptor. In this role, termed the MuSK-BMP pathway, MuSK binds BMPs 2, 4, and 7 via its Ig3 domain, as well as the type I BMP receptors BMPR1a and b (also termed Alk3 and Alk6, respectively). Cultured myoblasts and myotubes expressing MuSK show increased BMP signaling and distinctive transcriptomic responses compared to their MuSK-null counterparts (Yilmaz et al., 2016). To probe the function of the MuSK-BMP pathway more precisely, we generated mice lacking the BMP-binding MuSK Ig3 domain, “ΔIg3-MuSK”. Homozygous ΔIg3-MuSK mice are viable, born at normal ratios and survive at least 24 months (the oldest age examined). Moreover, agrin-induced, MuSK-mediated AChR clustering is normal in cultured ΔIg3-MuSK cells. At 3 months of age, the mice are similar to WT in weight and grip strength. Cells derived from these mice exhibit attenuated BMP-induced Smad1/5/8 phosphorylation and reduced levels of MuSK-dependent BMP-induced transcripts (Jaime et al., 2022, 2023). We have also used this model to show that the MuSK-BMP pathway acts in a cell autonomous manner in muscle stem (satellite) cells to regulate their quiescence and activation (Madigan et al., 2023), and that MuSK localized extrasynaptically in slow muscle is important for maintaining slow (but not fast) myofiber size via the Akt-mTOR pathway (Jaime et al., 2022, 2023). However, the function of the MuSK-BMP pathway at the NMJ is unknown.

Here, we investigated the structural and functional role of the MuSK-BMP pathway at the NMJ using the ΔIg3-MuSK mouse model. NMJs in ΔIg3-MuSK mice are fragmented throughout the lifespan, but anatomical innervation is preserved. Spontaneous and nerve-evoked ACh release as well as postsynaptic AChR density and currents in ΔIg3-MuSK NMJs are comparable to WT. In contrast, single-fiber electromyography (SFEMG) revealed MFAP jitter and blocking, indicating deficits in the ability of the nerve-evoked endplate potentials to induce muscle excitation. The amplitude of nerve-evoked compound MAPs (CMAPs) and muscle torque were also reduced. Finally, the level of Nav1.4 was reduced at ΔIg3-MuSK NMJs. We propose that MuSK plays two distinct roles at the NMJ: as an agrin-LRP4 receptor necessary for establishing and maintaining cholinergic signaling, and as a BMP co-receptor required for maintaining the structural integrity of the postsynaptic apparatus, Nav1.4 density, nerve-evoked muscle excitability and force production.

## Materials and methods

### Animals

All animal protocols were performed in compliance with regulations set by and with approval of the Brown Institutional Animal Care and Use Committee. ΔIg3-MuSK mice were created as described previously (Jaime et al., 2022) and maintained on the C57BL/6 background.

Transgenic Thy1-YFP mice (Feng et al., 2000), provided by Gregorio Valdez, were crossed with ΔIg3-MuSK and WT mice from the Fallon colony to generate animals with fluorescently-labeled motor axons.

### Immunohistochemistry

For staining of whole-mount muscle preparations, mice were sacrificed via CO_2_ inhalation followed by cervical dislocation or cardiac puncture to preserve the integrity of the neck muscles as needed. For most experiments in which we visualized NMJs, we used the sternomastoid, a flat, thin, and easily accessible muscle that is particularly suitable for NMJ morphology studies both in- and ex-vivo (Lichtman et al., 1987; Balice-Gordon and Lichtman, 1990; Bruneau and Akaaboune, 2006). Muscles were collected and pinned at resting length and connective tissue was removed. Muscles were then fixed in 4% paraformaldehyde for 20 minutes, fileted into bundles, washed with phosphate-buffered saline (PBS) 3×10 min, and labeled with tetramethylrhodamine-, Alexa Fluor 488-, or Alexa Fluor 647-conjugated ⍺-bungarotoxin (1:40, Invitrogen T1175, B13422, or B35450) for 15 minutes at room temperature to visualize AChRs. After washing, muscles were incubated in methanol at −20° C for 5 minutes, then washed again. Tissue was blocked for 1 hour in 0.2% Triton X-100, 2.0% bovine serum albumin (BSA) in PBS, then incubated with primary antibodies overnight with gentle agitation at 4°C. The next day, the muscles were washed 3 times for 10 minutes, incubated in AlexaFluor goat anti-rabbit 488 or 555 (1:200, Invitrogen A11008 or A21428) for 4 hours, then washed again before teasing into smaller bundles of muscle fibers and mounting in Vectashield mounting medium with DAPI (Vector, H-1200-10). Primary antibodies used were rabbit anti-neurofilament (1:2000, Sigma-Aldrich AB1987) to visualize axons, rabbit anti-VAChT (1:500, Synaptic Systems 139103) or rabbit anti-synaptophysin (Chemicon, discontinued) to visualize nerve terminals, mouse anti-Nav1.4 to visualize sodium channels (NeuroMab/Antibodies Inc N255/38), and rabbit anti-S100b (neat/ready-to-use, DAKO GA50461-2) to visualize terminal Schwann cells.

Nav1.4 staining was conducted in muscle cross-sections as previously described (Zhang et al., 2021) with slight modifications. Briefly, muscles were harvested and flash-frozen in liquid-nitrogen-frozen isopentane and embedded in optimum cutting temperature media. 10µm fresh-frozen sections were fixed with 4% PFA, washed, permeabilized for 10min in in 0.5% Triton X-100 in PBS. After blocking for one hour with 5% normal goat serum and 2% Triton X-100 in PBS, the primary antibody (1:1000, NeuroMab/Antibodies Inc N255/38) was applied in blocking buffer overnight at 4° C. After washing, secondary antibody (Alexafluor 594 Goat anti-Mouse IgG2a, Invitrogen, A-21135) and ⍺-bungarotoxin (item information above) were applied at 1:1000 for 1 hour. Slides were washed once more before mounting.

### Microscopy

Slides were blinded before imaging. Images were obtained using a Zeiss LSM 800 confocal microscope using either a 40x or 63x oil objective, and Zeiss Zen Blue software. Optical sections were taken at 2 µM intervals for 40x images, or 0.31 µM for detailed 63x images. For experiments examining Schwann cell numbers and Nav1.4 distribution in whole mounts, an Olympus FV3000 confocal laser scanning microscope with 60x oil objective was used. For quantitative immunofluorescence analysis of Nav1.4 in sections, images were taken using a Nikon ECLIPSE Ti2-E microscope. Images were collected at 20x magnification in a single session, using the same exposure for all slides.

### Image analysis

Z-stacks were collapsed into maximum intensity projections using ImageJ software. Postsynaptic fragmentation, or the number of segments of AChR, along with Schwann cell number were counted manually. When counting terminal Schwan cells, only Schwann cell bodies that overlapped the postsynaptic apparatus were counted, and Schwann cells associated only with the axon were excluded as S100β stains all Schwann cells.

We used a slightly modified version of aNMJ-morph to quantify other features of NMJ structure including areas of the pre- and post-synaptic apparatus and overall endplate region, and endplate compactness (Jones et al., 2016; Minty et al., 2020). Briefly, the macro separates the projection into two images, one per channel. The user (blinded to genotype) is instructed to threshold each channel, then “clean” or erase positive staining outside of the NMJ in each channel. The measurements are completed automatically by the macro. Because the aNMJ-morph macro was initially validated with 60x images, and images analyzed in this study were taken at 40x with larger intervals between optical sections, the measurements taken by aNMJ-morph that relied on ImageJ segmentation algorithms were not used.

Nav1.4 staining was quantified by tracing NMJs or similarly sized sarcolemmal areas to define regions of interest, then measuring mean intensity of Nav1.4 staining per region of interest. Background intensity was acquired from the center of the same myofiber. The mean intensity of Nav1.4 at neuromuscular junctions, minus the background, of WT and ΔIg3-MuSK TA muscles were then compared.

### Ex-vivo electrophysiology

Ex-vivo NMJ electrophysiology was conducted as previously described (Wang et al., 2004; Chugh et al., 2020). Briefly, the TA muscle was dissected, fileted and unfolded to create a flat surface. The muscle was then pinned and perfused with Ringer solution (physiological Ca^2+^ at 20-22° C, in 95% O_2_ and 5% CO_2_, see (Wang et al., 2004) for further details), then stained with 10uM 4-Di-2ASP and an epifluorescence microscope (Leica DMR) was used to visualize nerve terminals and muscle fibers. The fibers were then impaled within 100µM of the NMJ and crushed far from the motor endplate on either side to prevent contraction and movement. Under two-electrode voltage clamp at −45 mV, spontaneous miniature endplate current (mEPC) and evoked endplate current (EPC) amplitudes were recorded. For evoked endplate currents, a train of 10 stimulations at 50Hz was delivered to determine the level of decrement between the first and 10^th^ stimulation. Quantal content was calculated as the amplitude of a synapse’s EPC divided by the average mEPC amplitude.

### In-vivo Electrophysiology

Compound muscle action potentials (CMAP) were measured from the right hindlimb of WT and ΔIg3-MuSK mice as previously described (Arnold et al., 2015; Sheth et al., 2018). Briefly, mice were anesthetized with isoflurane (3-5% for induction and 1-2% for maintenance) delivered in compressed room air. The right hindlimb was shaved to allow for proper electrode contact. An active ring electrode was placed over the gastrocnemius muscle and a reference ring electrode was placed over the metatarsals of the right hindpaw (Alpine Biomed). Ring electrodes were coated with electrode gel (Spectra 360; Parker Laboratories) to increase contact with skin. A ground electrode was placed on the tail (Carefusion). One 28-gauge monopolar needle electrode (Teca, Oxford Instruments Medical, NY) was placed on either side of the sciatic nerve. A portable electrodiagnostic system (Natus, Middleton, WI) was used to stimulate the sciatic nerve (0.1 ms pulse, 1–10 mA intensity). CMAP baseline-to-peak amplitudes were recorded following supramaximal stimulation. Repetitive nerve stimulation (RNS) was carried out as previously described (Padilla et al., 2021) using the same electrode placement as in CMAP measurements. Trains of 10 supramaximal stimuli at 10 and 50 Hz were delivered and % decrement was calculated as (difference in CMAP amplitude between the 1^st^ and 10^th^ stimuli, divided by the amplitude of the first response) x 100.

Single-fiber electromyography (SFEMG) was carried out in mice anesthetized as above using an electrodiagnostic system with Viking software (Natus Neurology Inc) as previously described (Chugh et al., 2020; Padilla et al., 2021). The sciatic nerve was stimulated at 10 Hz using two 28-gauge monopolar needle electrodes. A strip ground electrode was placed on the opposite foot, and the recording electrode was inserted into the gastrocnemius muscle longitudinally to record muscle fiber action potentials (MFAPs). To be considered a MFAP, evoked responses had to have baseline-to-negative peak rise time of <500μs, baseline-to-negative peak amplitude ≥ 200μV and demonstrate an all-or-none response with appropriate waveform shape. The standard deviation of the latency between stimulation and peak of action potential response was calculated for 50-100 consecutive discharges per fiber. On average, 6 unique fibers were used per animal. MFAPs with jitter < 4µs were excluded to minimize the possibility of recording jitter from fibers due to direct muscle stimulation. Blocking was quantified as present or absent for each synapse assessed. Stimulation intensity was adjusted to confirm MFAP blocking was not attributable to submaximal stimulation.

### Measurement of muscle contractile torque force

In vivo measurements of plantarflexion torque were made as described previously (Sheth et al., 2018) using a muscle contractility apparatus (Model 1300A, Aurora Scientific Inc, Canada). With the foot and tibia aligned at 90°, the right foot was taped to the apparatus force plate and the knee was clamped at the femoral condyles, taking care to avoid compressing nearby nerves. Two electrodes inserted over the tibial nerve were used to stimulate plantarflexion. Peak twitch force was determined using a 1-second train of stimuli at 5Hz and quantifying the maximal twitch response. Maximum tetanic torque force was determined similarly using a 1s stimulus at 150 Hz. Force measurements were normalized to animal mass. Percent torque fade was calculated by the change of torque during a 1-second train was delivered at 150 Hz.

### Statistical Analysis

For morphometry analyses, data for each measurement was analyzed using R statistical software (Team, 2021) and R packages “MASS” (Venables and Ripley, 2002) and “betareg” (Cribari-Neto and Zeileis, 2010). Sample sizes (number of NMJs, number of animals per age and sex) are reported in Extended Data (Tables) for each dataset. Because the measurements were best fit by varying, non-normal statistical distributions, and in order to account for potential interaction between sex and genotype, generalized linear models were used in place of ANOVAs (which are based on the normal distribution) to allow for parametric testing. Distributions were selected using an unbiased method, minimization of the AIC (Akaike information criterion) among normal, lognormal and gamma for continuous data, Poisson or negative binomial for discrete data, and for proportion data the beta distribution was used. Animals from each age group were used only once and morphological parameters were not compared across time. Violin plots were made using the R packages “ggplot2” (Wickham, 2016) (part of the “tidyverse” family of packages (Wickham et al., 2019), which were used to prepare data for plotting) and arranged using “cowplot” (Wilke, 2020).

Statistical testing (unpaired Student’s T-test) for electrophysiology, muscle contractility, Schwann cell count and Nav1.4 were conducted in Prism 9. Data are shown as median +/-interquartile range for NMJ morphometry and mean +/-standard deviation for all other data unless otherwise stated, and threshold for significance is p<0.05.

### Code and Data Accessibility

The R script used for analysis and generation of plots, as well as the raw data for morphometric analysis are available upon request from the authors.

## Results

### NMJs in mature ΔIg3-MuSK mice are fragmented but anatomical innervation is preserved

As a first step to characterize the role of MuSK-BMP signaling at the NMJ, we examined NMJ morphology in sternomastoid (STM) muscles of male WT and ΔIg3-MuSK mice at 3 months of age. The most striking structural difference was postsynaptic fragmentation of ΔIg3-MuSK NMJs (Fig. 1a, b). For example, ∼40% of WT, but only ∼16% of ΔIg3-MuSK NMJs, had ≤ 4 AChR fragments. On the other hand, 26% of ΔIg3-MuSK NMJs had had ≥ 9 fragments, compared to 12% in WT. Female ΔIg3-MuSK mice also exhibited postsynaptic fragmentation compared to their WT counterparts (Figure 1c). In agreement with our earlier findings in 3-month-old EDL and soleus muscle (Jaime et al., 2022), we observed neither complete nor partial NMJ denervation in ΔIg3-MuSK mice of either sex.

**Figure 1.**
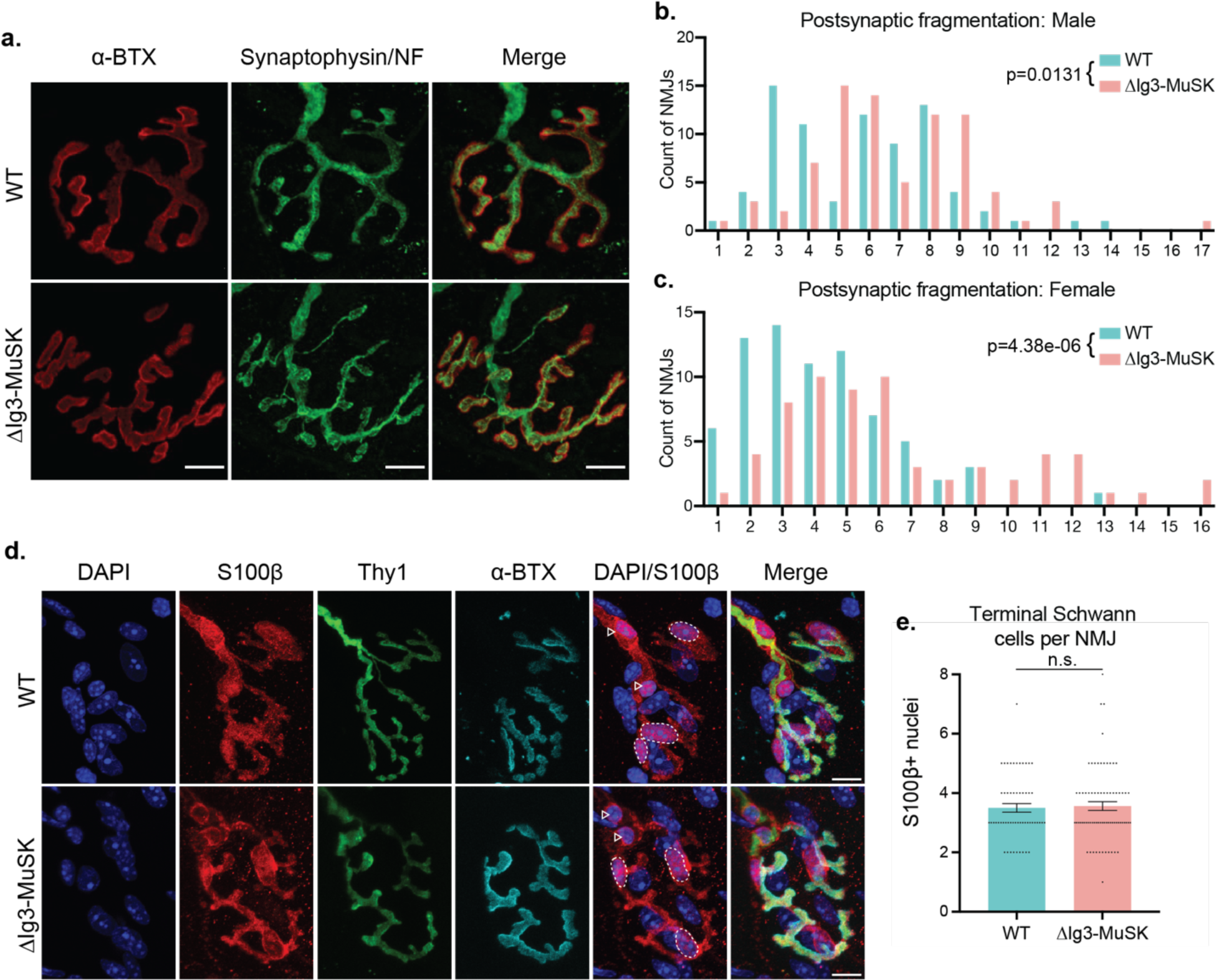
NMJs in young adult ΔIg3-MuSK mice are fragmented postsynaptically but maintain anatomical innervation and normal complement of terminal Schwann cells. **a.** WT and ΔIg3-MuSK NMJs with postsynaptic AChRs in red and nerve terminals and axons in green, bar = 10µm. Note the increase in postsynaptic fragmentation. **b.** Quantification of postsynaptic fragmentation in male WT and ΔIg3-MuSK mice. **c**. Quantification of postsynaptic fragmentation in female WT and ΔIg3-MuSK mice. (generalized linear models, full data and statistics presented in Tables 1-1 through 1-4, including median and interquartile range for each sex, n, test used.) **d.** Terminal Schwann cell bodies were identified by colocalization of S100β (red) and DAPI. Dotted outlines indicate terminal Schwann cell bodies (overlap of S100β and DAPI). Nerve terminals (Thy1-YFP) are shown in green, AChRs were visualized with ⍺-Bungarotoxin (cyan). Arrowheads indicate Schwann cells associated with the axonal input. **e.** The numbers of terminal Schwann cell bodies that overlapped or associated with AChRs were similar in WT and ΔIg3-MuSK NMJs (p = 0.772, unpaired T-test, n = 56 WT and 73 ΔIg3-MuSK NMJs).

### Terminal Schwann cells, overall synaptic morphological development and synapse elimination are not altered in ΔIg3-MuSK NMJs

Terminal Schwann cells play important roles at the NMJ (Feng and Ko, 2008; Kang et al., 2014; Lee et al., 2017); moreover, their numbers change during aging, which is also characterized by postsynaptic fragmentation (Fuertes-Alvarez and Izeta, 2021). We therefore asked whether the morphological changes noted above were accompanied by differences in the number and/or morphology of Schwann cells. As shown in Fig. 1d, e, WT and ΔIg3-MuSK NMJs had a similar number of terminal Schwann cells. Moreover, we did not observe changes in morphology, such as the presence of sprouts or blebs (Haizlip et al., 2015). Thus, the number and morphology of terminal Schwann cells is unaffected at the ΔIg3-MuSK NMJ.

### ΔIg3-MuSK NMJs exhibit increased postsynaptic fragmentation throughout the lifespan

We next sought to elucidate the contribution of the MuSK-BMP pathway to postnatal NMJ maturation and long-term stability by analyzing a set of WT and ΔIg3-MuSK NMJs from mice of both sexes across the lifespan (n=1619 NMJs). The murine NMJ undergoes extensive remodeling in the first weeks of postnatal life, progressing from an immature, polyinnervated and plaque-like configuration to a pretzel-like structure with a single axonal input (Figure 2a, (Balice-Gordon, 1997; Sanes and Lichtman, 2001; Shi et al., 2012). In both WT and ΔIg3-MuSK animals, NMJs at P14 had an overall small and compact appearance, with few if any branches or fragments (Fig 2b, c). From P21 through 3 months, NMJs in both genotypes grew in overall area and assumed a highly branched appearance. However, ΔIg3-MuSK NMJs had a higher count of postsynaptic fragments as early as P21. At P14 the middle 50% of the NMJs ranked from most to least fragments in both genotypes were fully continuous (1 fragment); by P21 WT and ΔIg3-MuSK NMJs had a median count of 2 and 3 postsynaptic fragments, respectively. This increase in fragmentation persisted through 24 months, the oldest age analyzed (Figure 2b-d, Tables 2-1 and 2-2). Importantly, we observed neither evidence of polyneuronal innervation nor partial or complete denervation in ΔIg3-MuSK compared to WT NMJs at any age.

**Figure 2.**
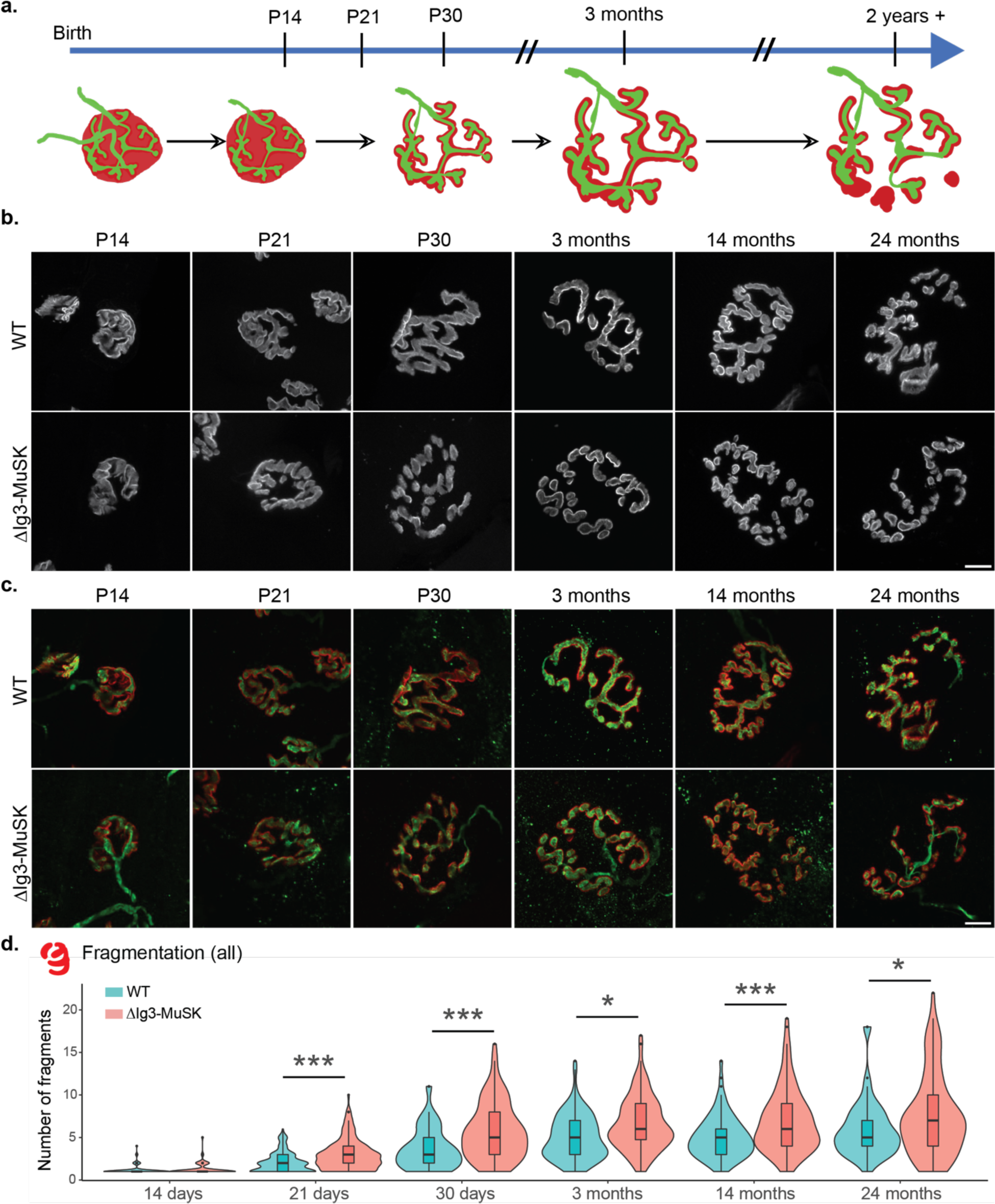
ΔIg3-MuSK NMJs exhibit increased postsynaptic fragmentation without anatomical denervation throughout the lifespan. **a.** Schematic of structural changes observed during WT NMJ development and aging. Note perforation of postsynaptic apparatus between birth and P30 and further postsynaptic fragmentation during aging. **b.** Postsynaptic structure (α-bungarotoxin only) of NMJs from sternomastoid of P14, P21, P30, and 3-, 14-, and 24-month old WT and ΔIg3-MuSK mice, **c.** merged images of pre- and post-synaptic apparatus of NMJs in (b). Postsynaptic apparatus (α-bungarotoxin, red); presynaptic (neurofilament with VAChT or synaptophysin; all green). Note increased fragmentation of postsynaptic elements and their persistent co-localization with presynaptic elements. **d.** Quantification of ΔIg3-MuSK NMJ fragmentation throughout the lifespan. (* p<0.05, ** p<0.01, *** p<0.001, generalized linear models, full statistics presented in Tables 2-1 and 2-2, including median and interquartile range for each measurement, n for each age and sex, tests used, and results of statistical testing.)

We next used the aNMJ-morph macro (Minty et al., 2020) to measure the size of the presynaptic and postsynaptic elements of WT and ΔIg3-MuSK NMJs at P14, P21, P30, 14 months, and 24 months. We conducted these measurements using both the combined sex dataset (Figure 2-1, Tables 2-1 and 2-2) and for each sex separately (Figure 2-2, Tables 1-1 through 1-4). At some ages we observed differences in AChR area and compactness, but unlike the fragmentation observation these changes were not consistent when compared across age and/or sex (Figure 2-2b-e, g-j). Importantly, fragmentation was observed in both sexes and all ages ≥ P21 (Figure 2-2, a, f).

### Postsynaptic fragmentation of ΔIg3-MuSK NMJs is observed in multiple muscle types

Different muscle types have characteristic NMJ morphologies and dynamics (Lømo and Waerhaug, 1985; Valdez et al., 2012). To assess whether the MuSK-BMP pathway plays a role in NMJ structure in both fast and slow muscle, we extended our morphometric analysis to NMJs in two hindlimb muscles - the fast extensor digitorum longus (EDL) and the slow soleus (SOL). We observed postsynaptic fragmentation of ΔIg3-MuSK NMJs in the EDL muscle, and a trend toward fragmentation in the soleus (p=0.0525, Figure 3a-c, Tables 3-1, 3-2). Interestingly, nerve terminal caliber was larger in the ΔIg3-MuSK soleus, but not the EDL (Fig 3d, e). Notably, NMJ size was also decreased in the soleus (Figure 3-1e, f, Tables 3-1, 3-2). We previously showed that myofiber diameter in the soleus, but not the fast TA, is reduced in ΔIg3-MuSK mice (Jaime et al., 2022, 2023). Thus, the MuSK-BMP pathway plays a role in maintaining NMJ structure in both fast and slow muscles. However, ΔIg3-MuSK NMJs in slow muscle show an additional phenotype where nerve terminal caliber is increased (Figure 3d-e), but the overall size of the nerve terminal and postsynaptic elements is smaller (see Discussion).

**Figure 3.**
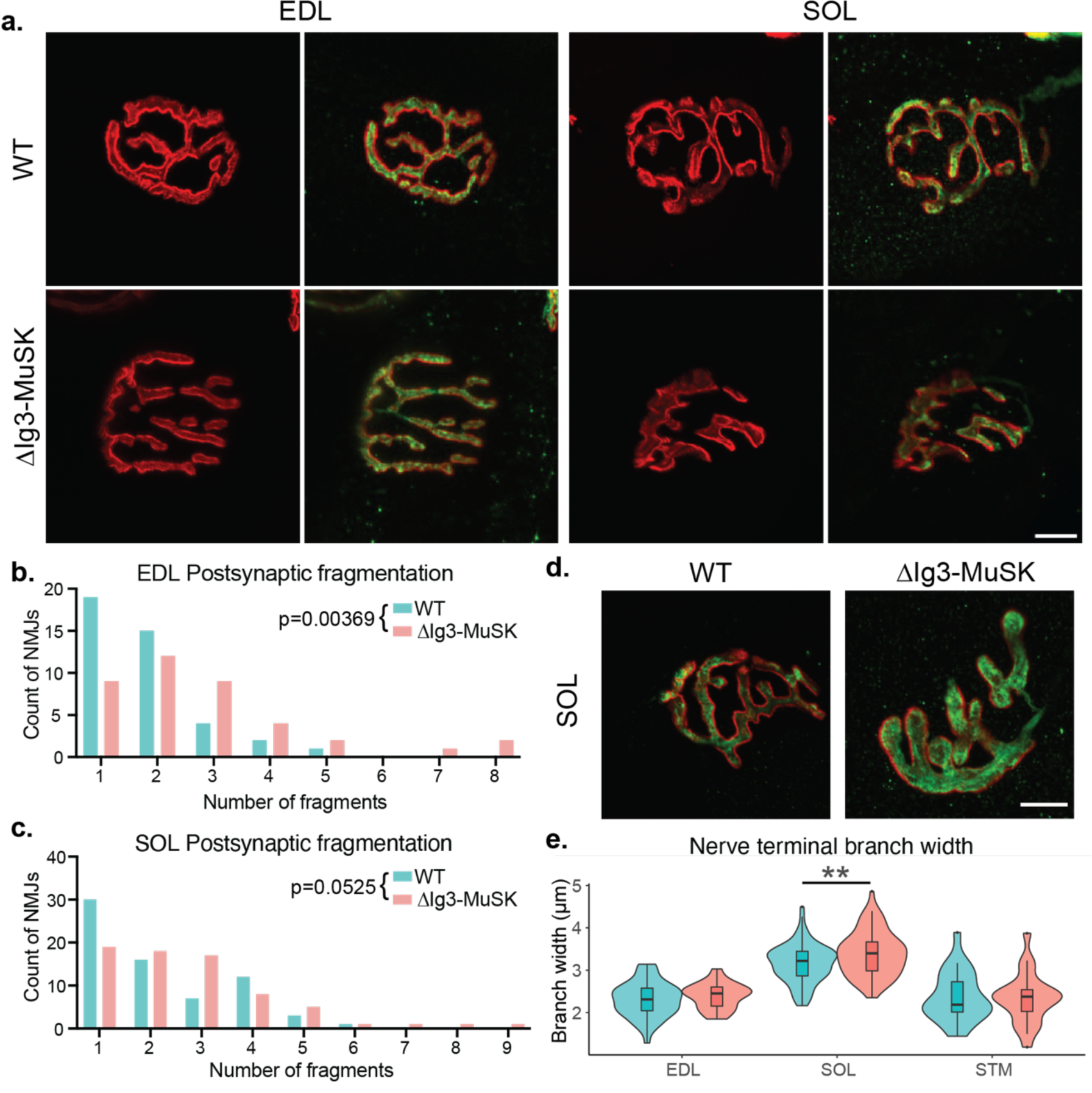
Postsynaptic fragmentation in ΔIg3-MuSK hindlimb muscles. **a.** WT and ΔIg3-MuSK EDL and soleus muscles were stained for pre-(green)and post-synaptic (red) elements for morphometric analysis. Both muscles exhibited postsynaptic fragmentation but showed no signs of denervation. **b**. Quantification of postsynaptic fragmentation observed in EDL. **c.** Quantification of postsynaptic fragmentation observed in SOL. **d.** Representative images illustrating wider nerve terminal branches in ΔIg3-MuSK Soleus. **e.** Quantification of average branch width in EDL, SOL, and STM NMJs from ΔIg3-MuSK and WT animals. Note increased branch width in SOL. (* p<0.05, ** p<0.01, *** p<0.001, generalized linear models, full statistics presented in Tables 3-1 and 3-2, including median and interquartile range for each measurement, n for each age and sex, tests used, and results of statistical testing.)

### Cholinergic function is preserved at ΔIg3-MuSK NMJs

We next assessed the role of the MuSK-BMP pathway in information transfer at the NMJ. Broadly considered, the function of the NMJ is to generate a MFAP in response to a nerve action potential. This process entails nerve-evoked cholinergic transmission to generate the endplate potential, and the activation of postsynaptic voltage-gated sodium channels (Nav1.4) to trigger the MFAP, or ‘excitability’. To characterize cholinergic signaling, we carried out ex-vivo measurements in the tibialis anterior under voltage clamp to quantify spontaneous miniature endplate currents (mEPCs), nerve-evoked endplate currents (EPCs), quantal content, and synaptic plasticity (depression/facilitation) in response to repetitive stimulation. All of these measures were comparable between ΔIg3-MuSK and WT synapses (Fig 4. a-d) (mEPC p=0.14, EPC p=0.74, quantal content p=0.61, depression/facilitation p=0.21; WT n=8, ΔIg3-MuSK n=4), 15-25 EPC/mouse). Importantly, the comparable size of the mEPCs and EPCs in the two genotypes provides direct evidence that postsynaptic AChR density is equivalent in ΔIg3-MuSK and WT NMJs. Further, the level of cholinergic signaling overall is comparable in both genotypes. Finally, the finding that quantal content and synaptic plasticity are equivalent at ΔIg3-MuSK and WT NMJs demonstrate that the number of vesicles released and the probability of their release are not affected by the ΔIg3-MuSK mutation. Taken together, these measures establish that the core elements of cholinergic signaling are preserved at the ΔIg3-MuSK NMJ.

**Figure 4.**
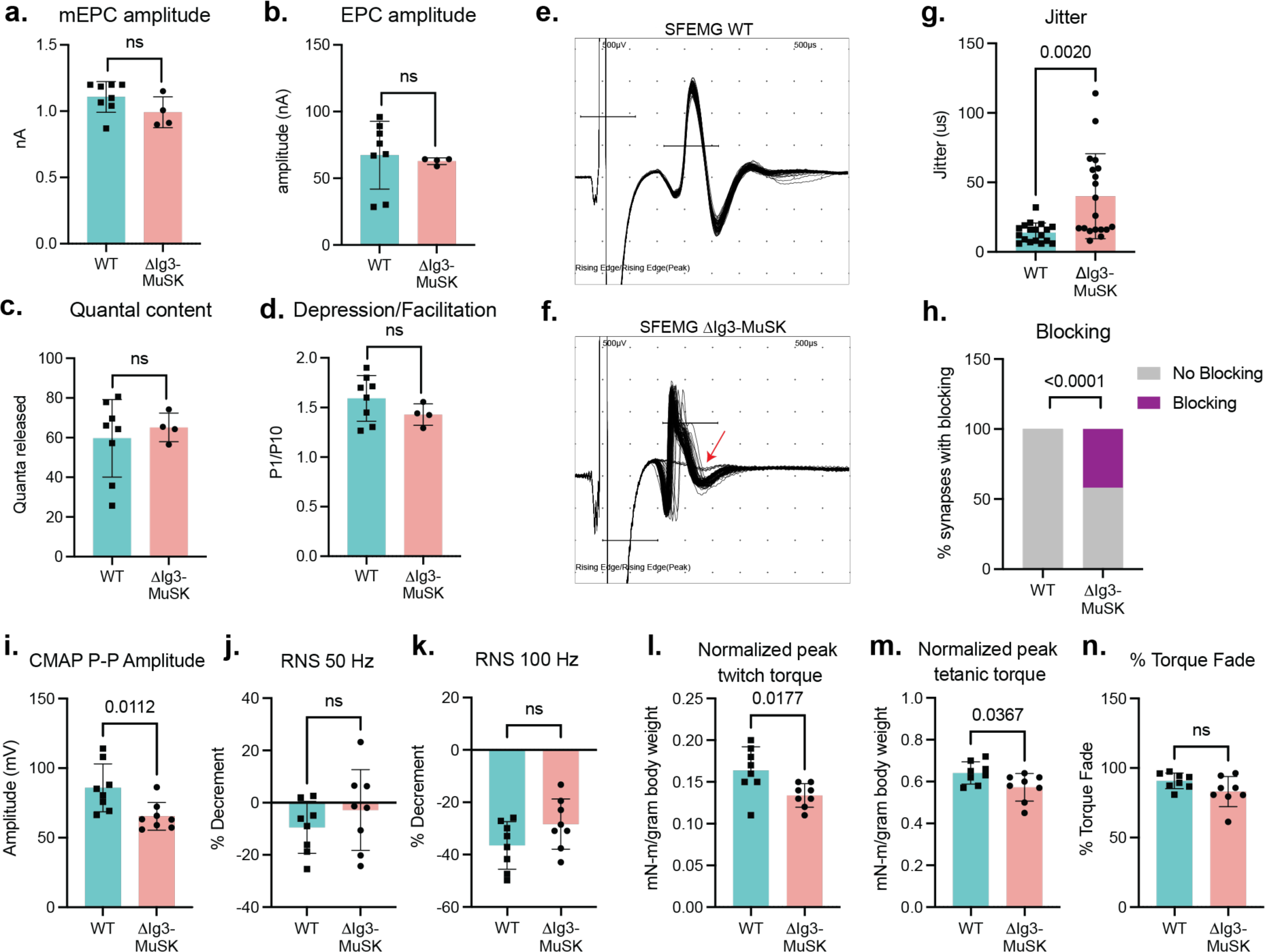
Impaired NMJ function in 6-month old ΔIg3-MuSK muscle. **a-d:** Cholinergic signaling. Ex-vivo measurements of tibialis anterior (**a.**) NMJ spontaneous miniature endplate current (mEPC) amplitude, (**b.**) nerve**-**evoked endplate current (EPC) amplitude, (**c.**) quantal content, (**d.**) and depression/facilitation in response to trains of stimulation. All four parameters were indistinguishable between WT and ΔIg3-MuSK muscle. (n: a-d 4 mice/genotype (b: 2 WT)), 18-26 mEPCs, EPCs, or trains of stimulation**). e, f:** Representative waveforms of single fiber electromyography (SFEMG) recordings of muscle fiber action potentials (MFAPs) in WT (**e.**) and ΔIg3-MuSK (**f.**) gastrocnemius. Note the inconsistent latency of MFAPs and blocking in the ΔIg3-MuSK muscle (red arrow). Both jitter and blocking (quantified in **g** and **h.,** respectively) were significantly increased in ΔIg3-MuSK (n = 3 mice/genotype, 5-7 trains of 50-100 stimuli per mouse; g. Student’s T-test, h. Chi-squared test.). **i.** Compound muscle action potential (CMAP) peak-to-peak amplitude is decreased in ΔIg3-MuSK (n: 6 mice/genotype, 1 stimulus per mouse), but CMAP % decrement in response to repetitive nerve stimulation (RNS) at **(j.)** 50 and **(k.)** 100 hz remains normal (n: 6 mice per genotype, 1 train of stimulus at each frequency per mouse). **l.** ΔIg3-MuSK mice produce less plantarflexion twitch torque in response to a single stimulus than WT. **m.** ΔIg3-MuSK mice produce less plantarflexion tetanic torque in response to a 1-second train of nerve stimuli than WT. **n.** Reduction of tetanic torque during a 1-s train of stimuli in ΔIg3-MuSK is similar to WT. (n: 6 mice/genotype, 1 twitch and 1 train of tetanic stimulus per mouse, Student’s t-test).

### Nerve-induced muscle excitability is impaired in ΔIg3-MuSK mice

We next tested whether the MuSK-BMP pathway plays a role in postsynaptic excitability and generation of MFAPs. We recorded nerve-evoked MFAPs from individual muscle fibers using single-fiber electromyography (SFEMG; Fig. 4 e, f) and measured both jitter, the variation in latency of MFAPs following trains of stimuli delivered at 1 Hz, and blocking, the failure to generate an MFAP in response to stimulus. We observed striking phenotypes in both features. Muscle fibers in ΔIg3-MuSK mice exhibited an >80% increase in jitter (Fig. 4 f, g, p = 0.002, Student’s T-test). Further, over 40% of the NMJs assessed exhibited blocking (failure of transmission), where a nerve stimulus fails to evoke a MFAP (Fig. 4 f, h, p < 0.0001, Chi-squared test).

To gain insight into neuromuscular transmission at the level of the whole muscle, we recorded compound muscle action potentials (CMAPs) in response to a single stimulation of the tibial nerve as well as decrement of response to repetitive nerve stimulation (RNS). As shown in Fig. 4i-k, while decrement in CMAP amplitude in response to repetitive nerve stimulation was similar between genotypes, the overall peak-to-peak amplitude of CMAPs was reduced in ΔIg3-MuSK mice. Taken together, these results show that the ability of the end plate potential to evoke MAPs is compromised in ΔIg3-MuSK muscle.

### Nerve-evoked muscle force is reduced in ΔIg3-MuSK animals

We next assessed the functional impact of the defects in NMJ excitability on nerve-induced muscle contraction. We measured twitch and tetanic plantarflexion torque force produced in response to tibial nerve stimulation. As shown in Fig. 4 (l,m), both twitch and tetanic torque force were decreased in the ΔIg3-MuSK animals (p=0.0177 and 0.0367, respectively). These in vivo electrophysiology and muscle contractility deficits show that ΔIg3-MuSK NMJs exhibit defective neuromuscular transmission. Taken together, these findings indicate that the MuSK-BMP pathway is important for muscle fiber excitability and force production.

### Nav1.4 density is reduced at ΔIg3-MuSK NMJs

Reliable NMJ excitability requires the localization of high densities of Nav1.4 channels at the synapse (Wood and Slater, 2001; Schiaffino and Reggiani, 2011; Zhang et al., 2021). We used immunostaining to compare the levels of Nav1.4 at WT and ΔIg3-MuSK NMJs in both whole mount and cross section in 6-month-old mice. As expected, Nav1.4 intensity was highest along the edges of the AChR-positive areas of the endplate (Figure 5a). To quantify the relative density of Nav1.4 we stained cross-sections of the tibialis anterior muscle at 6 months and conducted quantitative fluorescence intensity analysis for Nav1.4. As shown in Fig. 5b, c, Nav1.4 fluorescence intensity was decreased in ΔIg3-MuSK compared to WT at the NMJ (decreased by ∼27%; p<0.0001). Notably, extrajunctional Nav1.4 levels were comparable between genotypes (Fig. 5d). Therefore, we conclude that the MuSK-BMP pathway regulates NMJ excitability by selectively regulating levels of Nav1.4 at the synapse.

**Figure 5.**
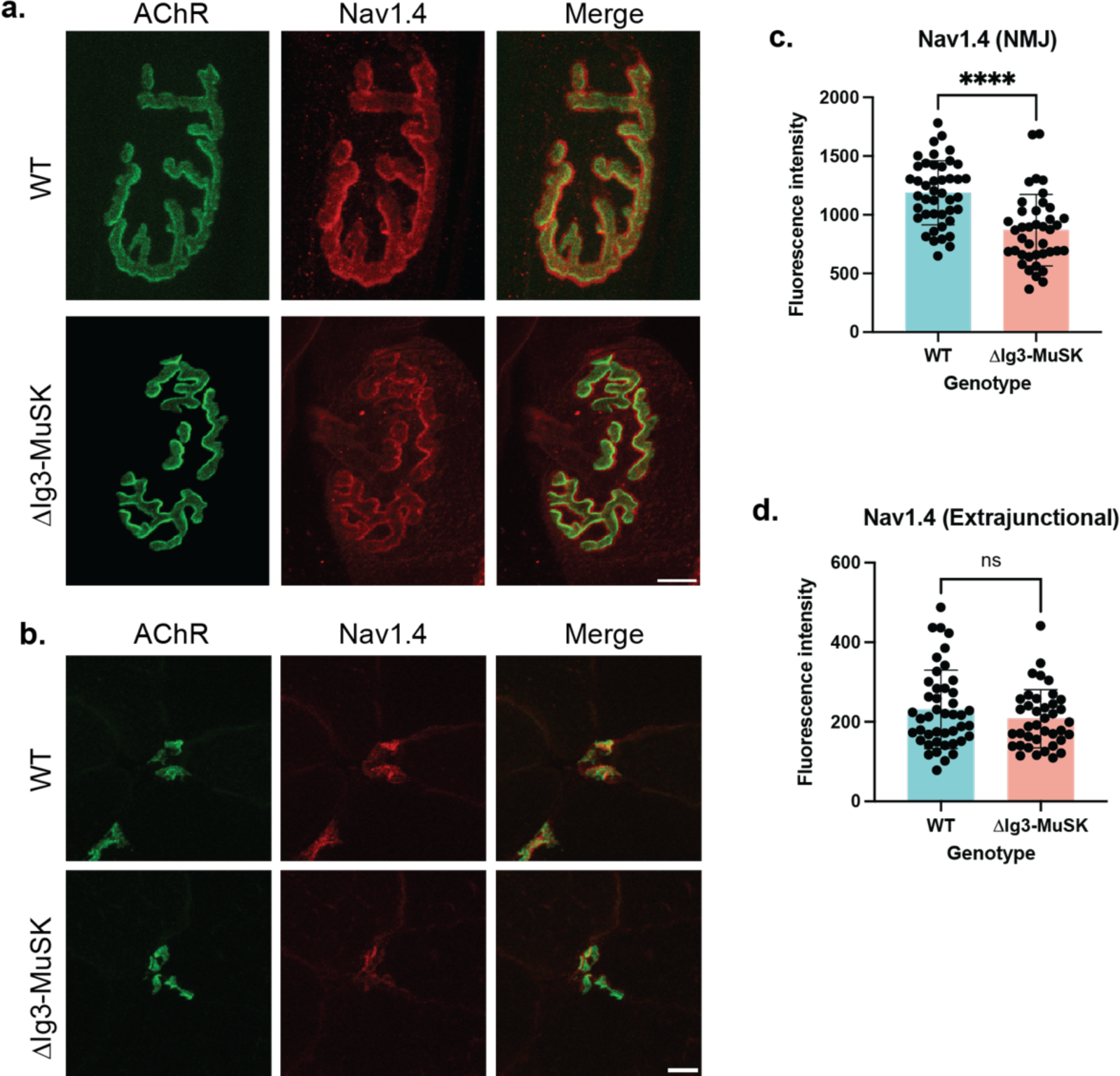
Decreased Nav1.4 levels at ΔIg3-MuSK NMJs. **a.** En face view of postsynaptic elements in the WT and ΔIg3-MuSK sternomastoid with AChRs in green and Nav1.4 in red. **b.** Cross-sectional staining of TA NMJs with postsynaptic AchRs in green and Nav1.4 in red. **c.** Quantification of Nav1.4 fluorescence intensity NMJs showed a statistically significant decrease in junctional Nav1.4 (p<0.0001, unpaired T-test, n= 43 NMJs from 4 WT mice, 40 NMJs from 4 ΔIg3-MuSK mice). **d.** Extrajunctional sarcolemmal Nav1.4 fluorescence intensity was comparable between WT and ΔIg3-MuSK (p= 0.22, unpaired T-test, n=44 regions of interest from 4 WT mice, 39 regions of interest from 4 ΔIg3-MuSK mice).

## Discussion

In this study we show that the MuSK-BMP pathway is important for NMJ structure and function. Taken together with previous work, we propose that MuSK plays two distinct roles at the NMJ: as an agrin-LPR4 receptor necessary for organizing the cholinergic signaling apparatus, and as a BMP co-receptor necessary for establishing normal NMJ structure, Nav1.4 density and NMJ excitability (Fig 6). Here we discuss these findings and their implications related to aging and diseases affecting the neuromuscular system.

**Figure 6.**
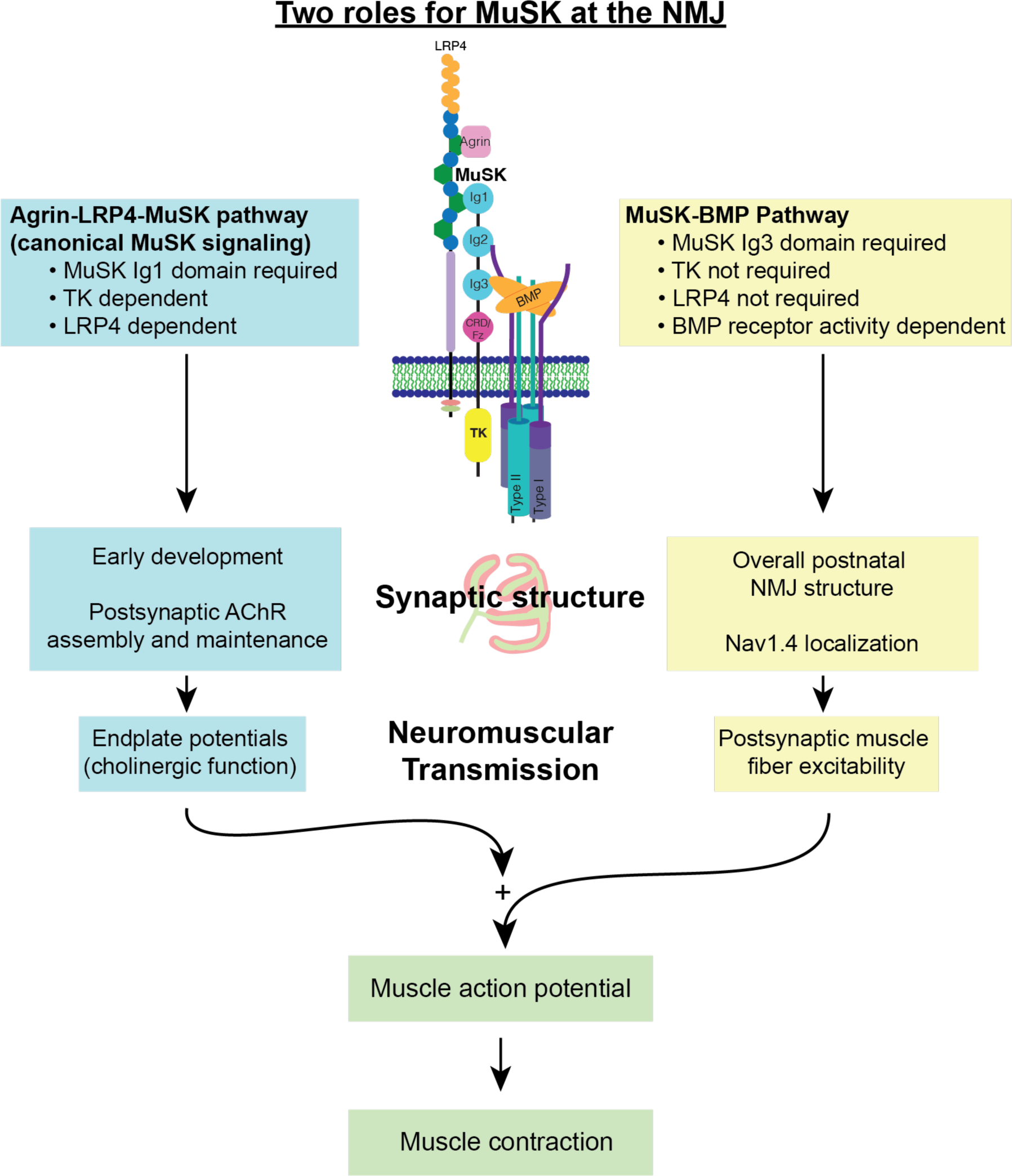
Two roles for MuSK at the NMJ. MuSK acts as both an agrin-LRP4 receptor and BMP co-receptor at the NMJ. In these two roles, MuSK regulates different elements of postsynaptic machinery and NMJ function. Both roles of MuSK are required for endplate potentials to be produced and be reliably amplified into action potentials, initiating a muscle contraction.

The MuSK-BMP pathway is necessary for maintaining NMJ integrity. The most prominent structural phenotype of ΔIg3-MuSK NMJs is fragmentation of the postsynaptic apparatus, which manifests as early as P21. Fragmentation remained elevated across the entire age range studied (up to 24 months; Fig. 2). This structural defect was not secondary to muscle damage (Li et al., 2011; Rudolf et al., 2014; Slater, 2020), since neither myofiber death nor regeneration is observed in ΔIg3-MuSK mice (Jaime et al., 2022). This fragmentation is also observed far earlier than in aging WT muscles, which show this phenotype only beginning around 18 months (Valdez et al., 2010; Fish and Fallon, 2020), raising the possibility that the MuSK-BMP pathway is one of mechanisms that is compromised during normal aging. Further, the MuSK-BMP pathway could be a general mechanism for maintaining NMJ structural integrity as we observe increased fragmentation in hindlimb muscles (Fig. 3). Interestingly, there was a soleus-selective increase in nerve terminal caliber in 3-month-old ΔIg3-MuSK animals that was independent of overall size of the synapse as measured by nerve terminal area and AChR area (which were smaller in ΔIg3-MuSK) or overall area of the endplate, which was unchanged.

Since MuSK is not expressed in motor neurons, this increase could reflect a MuSK-BMP dependent retrograde signal that regulates nerve terminal size. Alternatively, it could be the result of a homeostatic response to the reduced slow (but not fast) myofiber and NMJ size at 3 months of age ((Jaime et al., 2022, 2023), Figure 3-1). Taken together, these observations suggest that the MuSK-BMP pathway acts to maintain the structural integrity of the postsynaptic apparatus.

Our results establish that the MuSK-BMP pathway is necessary for reliable nerve-muscle communication. SFEMG revealed increased jitter and blocking in MFAPs generated in response to nerve stimulation, despite preservation of normal cholinergic signaling at the ΔIg3-MuSK NMJ. This excitability defect is the likely basis for the observed reduction in CMAPs and nerve-induced muscle force in the mutant muscle. The reduced excitability is likely due to the diminished localization of Nav1.4 at the synapse. Notably, the level of Nav1.4 in the non-synaptic sarcolemma is comparable in WT and ΔIg3-MuSK muscle. These observations are supported by a recent study where myofiber-specific knockout of ankyrins in muscle resulted in a complete loss of Nav1.4 at the NMJ, but not the sarcolemma. Further, in that study CMAP fatigue and reduced running activity were also observed (Zhang et al., 2021). We thus propose that the MuSK-BMP pathway functions at the level of the synapse to regulate muscle excitability via Nav1.4 localization.

MuSK’s canonical signaling pathway requiring its Ig1 and tyrosine kinase domains is crucial for the organization of cholinergic receptors at the NMJ and maintaining innervation (Yumoto et al., 2012; Burden et al., 2013; Tintignac et al., 2015). Our previous work showed that the canonical MuSK signaling pathway is intact in ΔIg3-MuSK cells and that MuSK-BMP signaling is neither activated by agrin nor requires MuSK tyrosine kinase function (Yilmaz et al., 2016; Jaime et al., 2022, 2023). Further, the ex-vivo electrophysiology experiments in the current work confirm that AChR density and cholinergic currents are normal at ΔIg3-MuSK NMJs while Nav1.4 localization and muscle excitability are compromised. Finally, it is noteworthy that full length agrin can bind BMP4 via follistatin-like domains in its N-terminal domain (Bányai et al., 2010) and can induce the clustering of Nav1.4 on cultured myotubes. In contrast, c-terminal fragments of agrin can activate LRP4-MuSK signaling and AChR aggregation but fail to induce Nav1.4 clustering (Sharp and Caldwell, 1996). These observations raise the possibility that agrin may participate, in a domain-specific manner, in both MuSK-BMP and MuSK-LRP4 signaling at the synapse.

Our findings also have implications for neuromuscular disease and aging. Auto antibodies to MuSK underlie a clinical presentation of Myasthenia Gravis (MG) distinct from AChR antibody-mediated MG, that is accompanied by muscle atrophy and can mimic early symptoms of amyotrophic lateral sclerosis (ALS) (Furuta et al., 2015; Huijbers et al., 2016). The best understood mechanism of MuSK-MG is disruption of agrin-LRP signaling (Huijbers et al., 2013; Fish and Fallon, 2020). It will be of interest to determine whether autoimmune disruption of MuSK-BMP signaling via antibody binding to the MuSK Ig3 domain might also contribute to MuSK MG, either through targeting the MuSK Ig3 domain or by antibodies that modulate MuSK binding to Type I BMP receptors, which does not require the Ig3 domain (Yilmaz et al., 2016; Fish and Fallon, 2020).

Finally, our findings provide a novel mechanistic framework for understanding age-related sarcopenia. In humans, the muscle weakness that characterizes sarcopenia progresses 2-5 fold faster than the decrement in muscle size (Mitchell et al., 2012), suggesting that the loss of muscle function could be due to defects in excitability independent of changes in anatomical innervation or cholinergic signaling. This model is supported by our recent findings in aged WT mice and rats, which showed reduced nerve-evoked MAPs and muscle force, but no defects in cholinergic signaling (Sheth et al., 2018; Chugh et al., 2020; Padilla et al., 2021). In aged humans, relatively small studies attempting to standardize clinical reference values have shown that SFEMG jitter increases with age (Bromberg et al., 1994; Balci et al., 2005). One small-scale study also showed an increase in jitter along the timeline from pre-sarcopenia to severe sarcopenia (Gilmore et al., 2017). However, there still has not been a comprehensive, large-scale SFEMG study on sarcopenic individuals (Tintignac et al., 2015). Taken together, these findings point to the NMJ as an important focus of sarcopenia pathology and suggest that the MuSK Ig3 domain, and potentially the MuSK-BMP pathway, could inform upon therapeutic targets for ameliorating this devastating condition in aging humans.

## Author contributions

LAF: Designed and interpreted all experiments, executed experiments in Figs 1-3, 5, wrote manuscript. MDE: executed and interpreted experiments (Figs. 1-3), contributed writing to paper. DJ: generated ΔIg3-MuSK mouse line. CX: executed experiment in Fig. 5e-f. KAR: conceptualized, executed, interpreted experiments in Fig. 4e-n. XW: executed experiments in Fig 4a-d. REF: collected data for Fig. 1h-i. MMR: conceptualized and interpreted experiments in Fig 4a-d. WDA: conceptualized, executed, and interpreted experiments in Fig 4e-n. JRF: Designed and interpreted experiments, wrote manuscript.

## Acknowledgements

We thank Beth McKechnie for excellent technical support, Gregorio Valdez for providing Thy1-YFP mice, Geoffrey Williams for his expertise in microscopy and generosity with his time, and Joseph Langan for his support with statistical analysis. LAF was supported by T32 MH20068 and a Carney Institute Graduate Award. MDE was supported by a Brown University UTRA. DJ was supported by 4R25GM083270 and 2T32AG041688, KAR was supported by 1F99AG079815-01 and the Ohio State University Presidential Fellowship, MMR and XW were supported by AR074985, KAW was supported by an Emerging Areas of New Science DEANS Award from the Division of Biology and Medicine at Brown University, the Cleft Palate Foundation, and R01 GM068118, JRF was supported by U01 NS064295, R41 AG073144, R21 NS112743, R21 AG073743, 1S10OD025181 and ALS Finding a Cure.

## Extended Data Figure Legends

**Figure 2-1.**
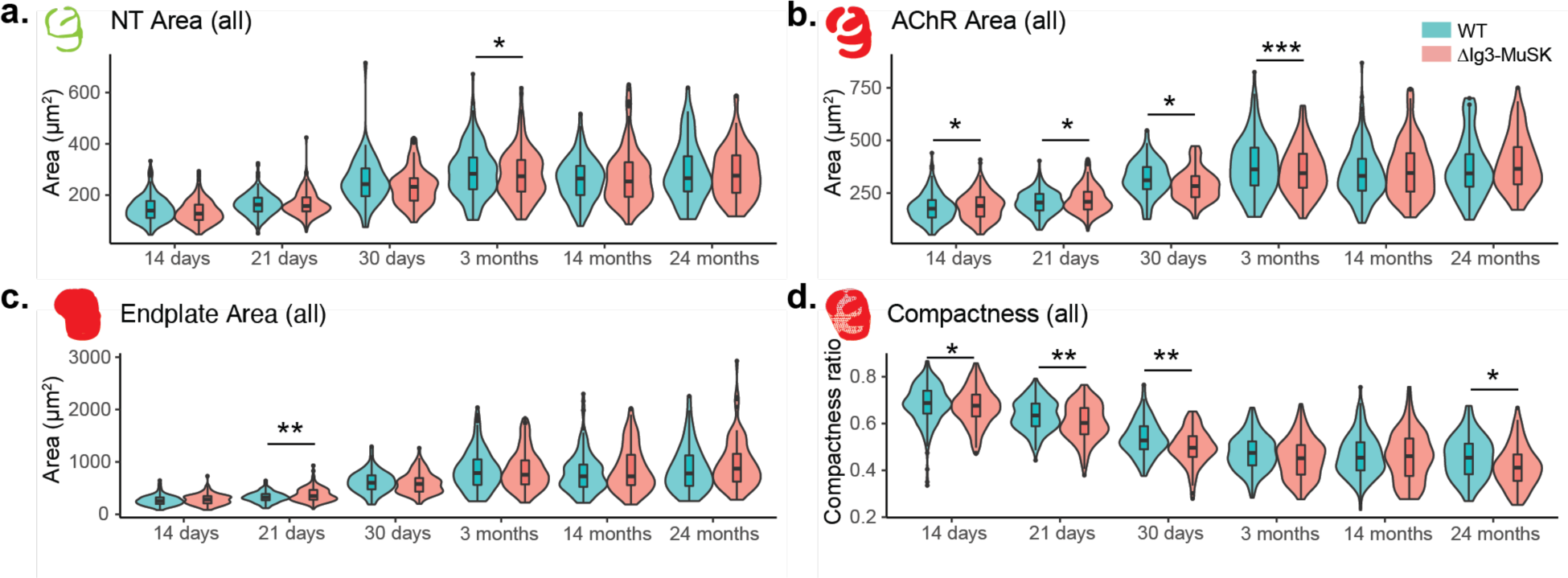
Quantification of nerve terminal area (**a.**), AChR area (**b.**), endplate area (**c.**), overall area of the endplate region, and (**d.**) compactness ratio of ΔIg3-MuSK NMJs throughout the lifespan. (* p<0.05, ** p<0.01, *** p<0.001, generalized linear models, complete statistics are presented in Tables 2-1 and 2-2, including median and interquartile range for each measurement, n for each age and sex, tests used.)

**Figure 2-2.**
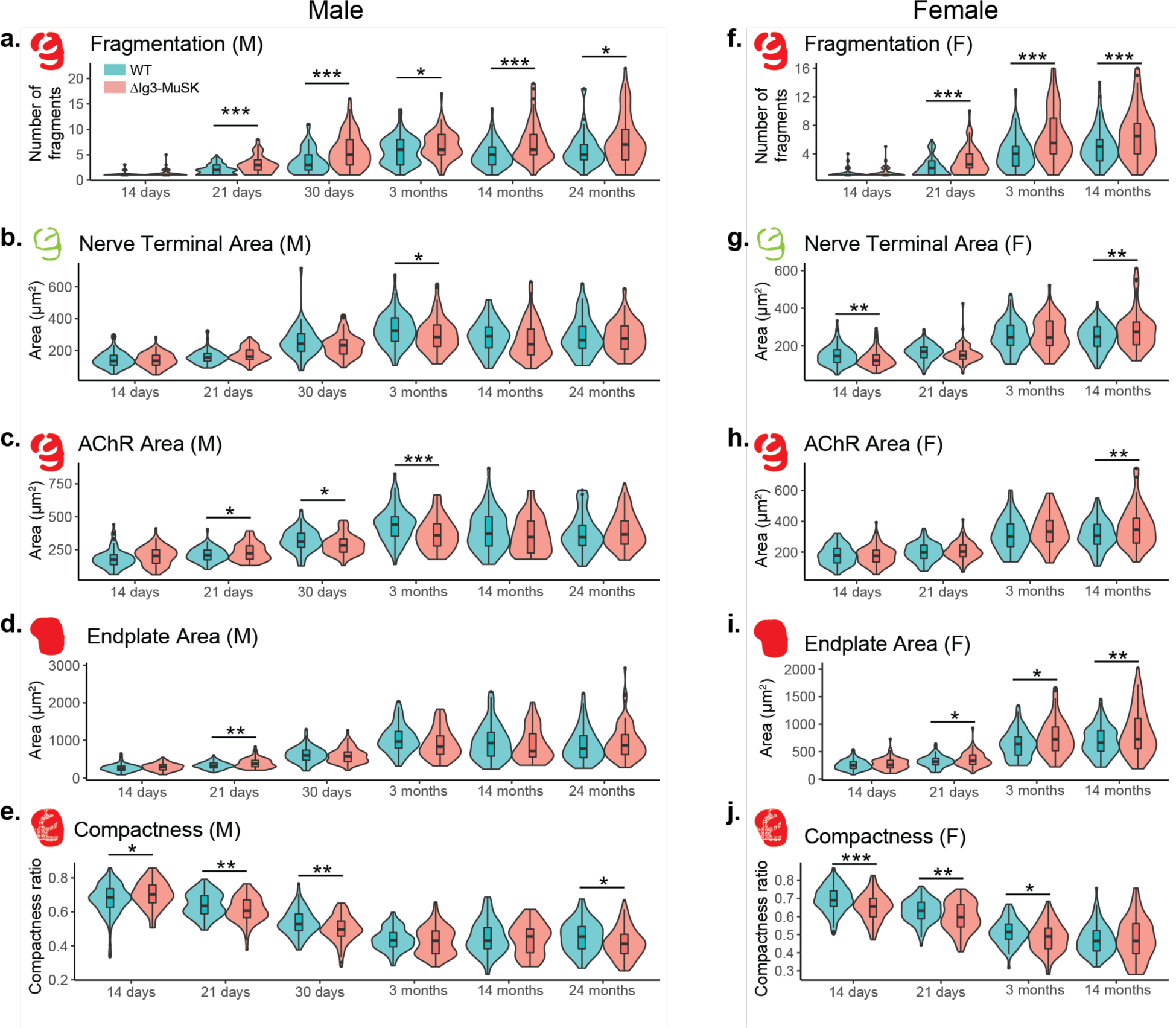
Fragmentation and other NMJ metrics measured across the lifespan for male and female ΔIg3-MuSK mice separately. Morphometric analysis of NMJs in males (**a-e**) and female (**f-j**) at the indicated ages. Fragmentation (**a, f)**, nerve terminal area (**b, g**), AChR area (**c, h**), endplate area (**d, i**), overall area of the endplate region, and (**e, j**) compactness ratio of male ΔIg3-MuSK NMJs throughout the lifespan. (* p<0.05, ** p<0.01, *** p<0.001, generalized linear models, full statistics are presented in Tables 1-1 and 1-2 (male), 1-3 and 1-4 (female), including median and interquartile range for each measurement, n for each age and sex, tests used, and results of statistical testing.)

**Figure 3-1.**
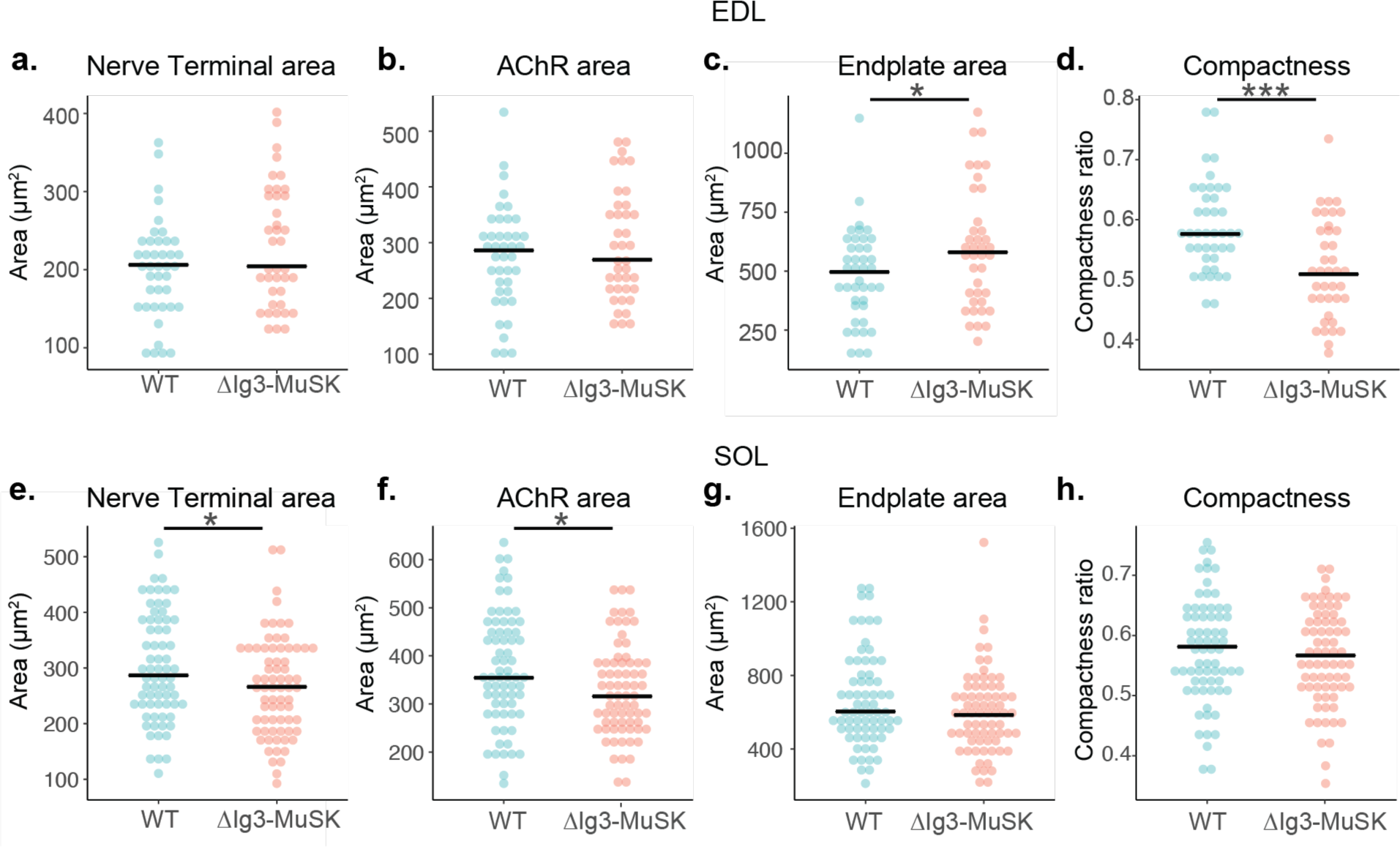
Additional NMJ morphometric data from ΔIg3-MuSK EDL and SOL. **a**. Morphometric analysis of NMJs in EDL (***a-d***) and soleus (**e-h**) at 3 months of age. Nerve terminal area (**a, e**), AChR area (**b, f**), endplate area (**c, g**), overall area of the endplate region, and (**d, h**) compactness ratio of male ΔIg3-MuSK NMJs at 3 months of age. (* p<0.05, ** p<0.01, *** p<0.001, generalized linear models, full statistics are presented in Tables 3-1 and 3-2).

## Extended Data Table Legends

**Table 1-1.** Male-only sternomastoid NMJ morphometry data across lifespan. Data for each timepoint studied presented as: Median value (interquartile range). Sample sizes (animals and synapses) indicated for each timepoint.

Extended data: Tables
| Age | P14 |  | P21 |  |
| --- | --- | --- | --- | --- |
| Genotype | WT | ΔIg3-MuSK | WT | ΔIg3-MuSK |
| Nerve Terminal Area ($\mu\text{m}^2$) | 134.374 (108.17, 169.624) | 135.632 (108.061, 170.442) | 155.226 (133.552, 181.631) | 162.096 (143.183, 204.451) |
| AChR Area ($\mu\text{m}^2$) | 174.074 (135.389, 211.237) | 202.719 (145.052, 250.194) | 207.802 (173.411, 247.451) | 222.007 (176.808, 278.327) |
| Endplate Area ($\mu\text{m}^2$) | 249.85 (200.152, 315.087) | 289.332 (213.942, 350.153) | 321.944 (269.604, 396.63) | 372.687 (294.528, 465.446) |
| Compactness | 0.685 (0.627, 0.737) | 0.703 (0.653, 0.759) | 0.635 (0.59, 0.696) | 0.607 (0.566, 0.67) |
| Fragmentation | 1 (1, 1) | 1 (1, 1) | 2 (1, 3) | 3 (2, 4) |
| N NMJs) | 103 | 77 | 85 | 82 |
| N (animals) | 6 M | 5 M | 6 M | 6 M |

| Age | P30 |  | 3 months |  |
| --- | --- | --- | --- | --- |
| Genotype | WT | ΔIg3-MuSK | WT | ΔIg3-MuSK |
| Nerve Terminal Area ($\mu\text{m}^2$) | 242.712 (194.96, 303.927) | 231.833 (177.882, 265.737) | 324.85 (256.83, 404.88) | 283.90 (224.97, 359.78) |
| AChR Area ($\mu\text{m}^2$) | 310.453 (268.522, 373.865) | 283.402 (230.952, 330.542) | 440.97 (351.43, 499.85) | 359.27 (278.03, 446.45) |
| Endplate Area ($\mu\text{m}^2$) | 599.541 (475.166, 742.781) | 579.201 (431.658, 690.198) | 965.78 (794.17, 1238.18) | 834.11 (624.07, 1117.29) |
| Compactness | 0.528 (0.49, 0.588) | 0.496 (0.457, 0.546) | 0.433 (0.395, 0.478) | 0.429 (0.353, 0.487) |
| Fragmentation | 3 (2, 5) | 5 (3, 8) | 6 (3, 8) | 6 (5, 9) |
| N (NMJs) | 61 | 89 | 77 | 80 |
| N (Animals) | 5 M | 7 M | 6 M | 6 M |
| Nerve terminal caliber (Fig. 3) | NA | NA | 2.187 (2.01, 2.726) | 2.375 (2.026, 2.541) |

**Table 1-1. Male-only sternomastoid NMJ morphometry data across lifespan**
| Age | 14 months |  | 2 Years |  |
| --- | --- | --- | --- | --- |
| Genotype | WT | Δlg3-MuSK | WT | Δlg3-MuSK |
| Nerve Terminal Area ($\mu\text{m}^2$) | 288.367 (216.764, 347.376) | 238.753 (173.655, 334.442) | 265.435, (214.067, 351.285) | 275.408, (208.403, 354.959) |
| AChR Area ($\mu\text{m}^2$) | 371.013 (277.396, 500.124) | 345.33 (224.661, 467.449) | 343.954, (280.852, 434.392) | 365.561, (291.658, 468.384) |
| Endplate Area ($\mu\text{m}^2$) | 926.124 (580.3, 1214.764) | 716.36 (556.243, 1176.327) | 778.891, (541.145, 1122.401) | 870.286, (623.292, 1152.434) |
| Compactness | 0.429 (0.384, 0.507) | 0.455 (0.359, 0.499) | 0.454 (0.383, 0.514) | 0.411 (0.354, 0.468) |
| Fragmentation | 5 (3, 6.5) | 6 (5, 9) | 5 (4, 7) | 7 (4, 10) |
| N (NMJs) | 63 | 53 | 53 | 86 |
| N (animals) | 5 M | 5 M | 5 M | 7 M |

**Table 1-2.** Male-only sternomastoid statistical results across lifespan. Best fit distribution used to fit GLM for each morphological measurement and associated p-values for genotype effect. GLMs fit by minimization of AIC. Significant p-values in bold.

| Measurement | Statistics | P14 | P21 | P30 | 3 months | 14 months | 2 years |
| --- | --- | --- | --- | --- | --- | --- | --- |
| **Nerve** | Distribution | gamma | gamma | gamma | gamma | gamma | gamma |
| **Terminal Area** | Genotype P Value | 0.692 | 0.0557 | 0.0652 | **0.041** | 0.23 | 0.867 |
| **AChR Area** | Distribution | gamma | gamma | gamma | gamma | gamma | gamma |
|  | Genotype P Value | 0.0623 | **0.0154** | **0.0164** | **0.0008** | 0.247 | 0.344 |
| **Endplate Area** | Distribution | gamma | gamma | gamma | gamma | gamma | gamma |
|  | Genotype P Value | 0.11 | **0.00112** | 0.637 | 0.0519 | 0.482 | 0.281 |
| **Compactness** | Distribution | beta | beta | beta | beta | beta | beta |
|  | Genotype P Value | **0.0219** | **0.00721** | **0.00192** | 0.335 | 0.548 | **0.0135** |
| **Fragmentation** | Distribution | poisson | poisson | negative binomial | negative binomial | negative binomial | negative binomial |
|  | Genotype P Value | 0.55 | **6.18e-05** | **1.89e-05** | **0.0131** | **8.49e-05** | **0.0236** |

**Table 1-3.**
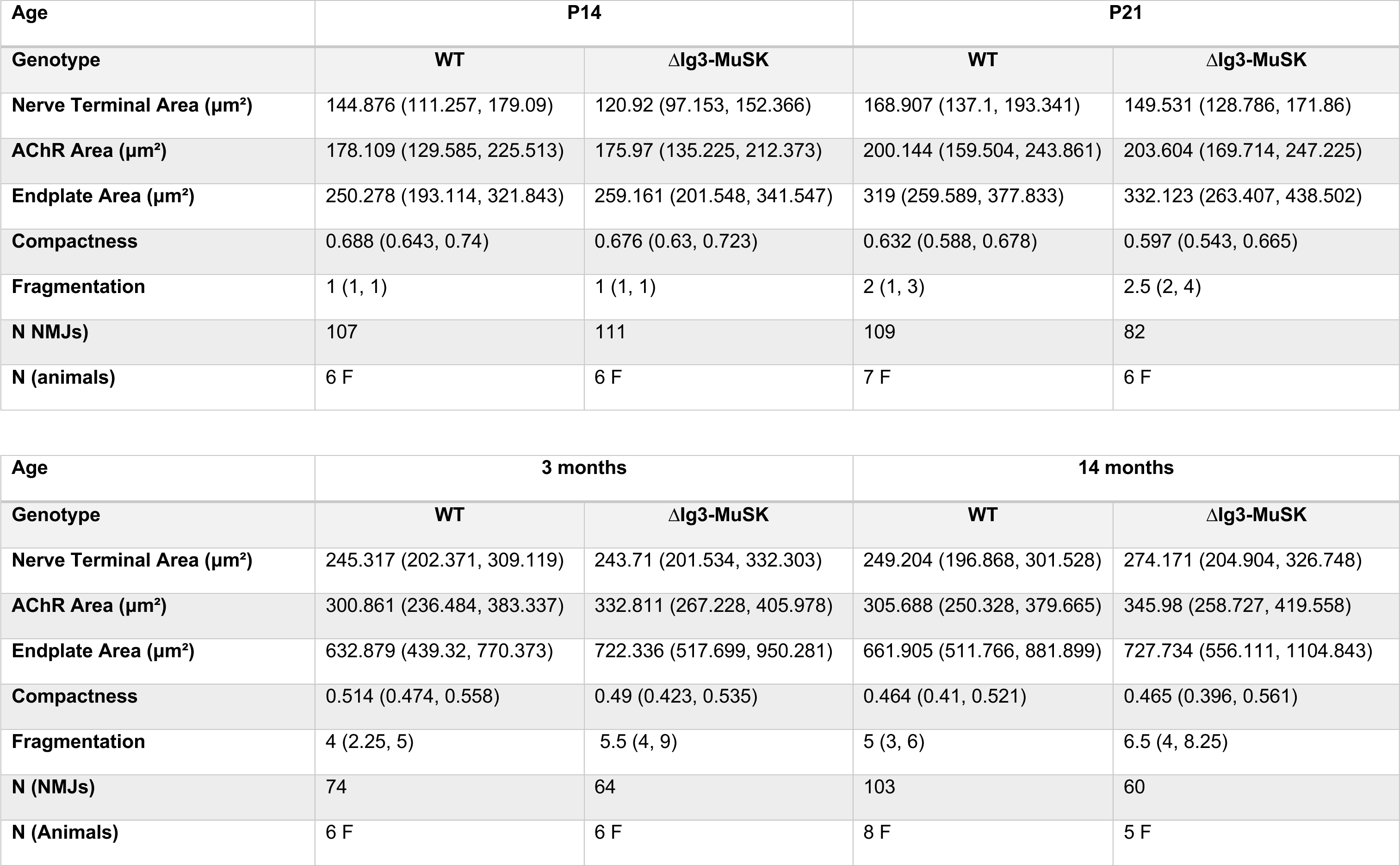
Female-only sternomastoid NMJ morphometry data across lifespan. Data for each timepoint studied presented as: Median value (interquartile range). Sample sizes (animals and synapses) indicated for each timepoint.

**Table 1-4.** Female sternomastoid statistical results across lifespan. Best fit distribution used to fit GLM for each morphological measurement and associated p-values for genotype effect. GLMs fit by minimization of AIC. Significant p-values in bold.

| Measurement | Statistics | P14 | P21 | 3 months | 14 months |
| --- | --- | --- | --- | --- | --- |
| **Nerve** | Distribution | gamma | gamma | gamma | gamma |
| **Terminal Area** | Genotype P Value | **0.00676** | 0.141 | 0.411 | **0.00739** |
| **AChR Area** | Distribution | gamma | lognormal | gamma | gamma |
|  | Genotype P Value | 0.705 | 0.44 | 0.12 | **0.00481** |
| **Endplate Area** | Distribution | gamma | gamma | gamma | gamma |
|  | Genotype P Value | 0.223 | **0.0383** | **0.0197** | **0.00607** |
| **Compactness** | Distribution | beta | beta | beta | beta |
|  | Genotype P Value | **9.87e-06** | **0.00248** | **0.0208** | 0.85155 |
| **Fragmentation** | Distribution | poisson | poisson | negative binomial | negative binomial |
|  | Genotype P Value | 0.6258 | **6.58e-05** | **4.38e-06** | **0.000643** |

**Table 2-1.**
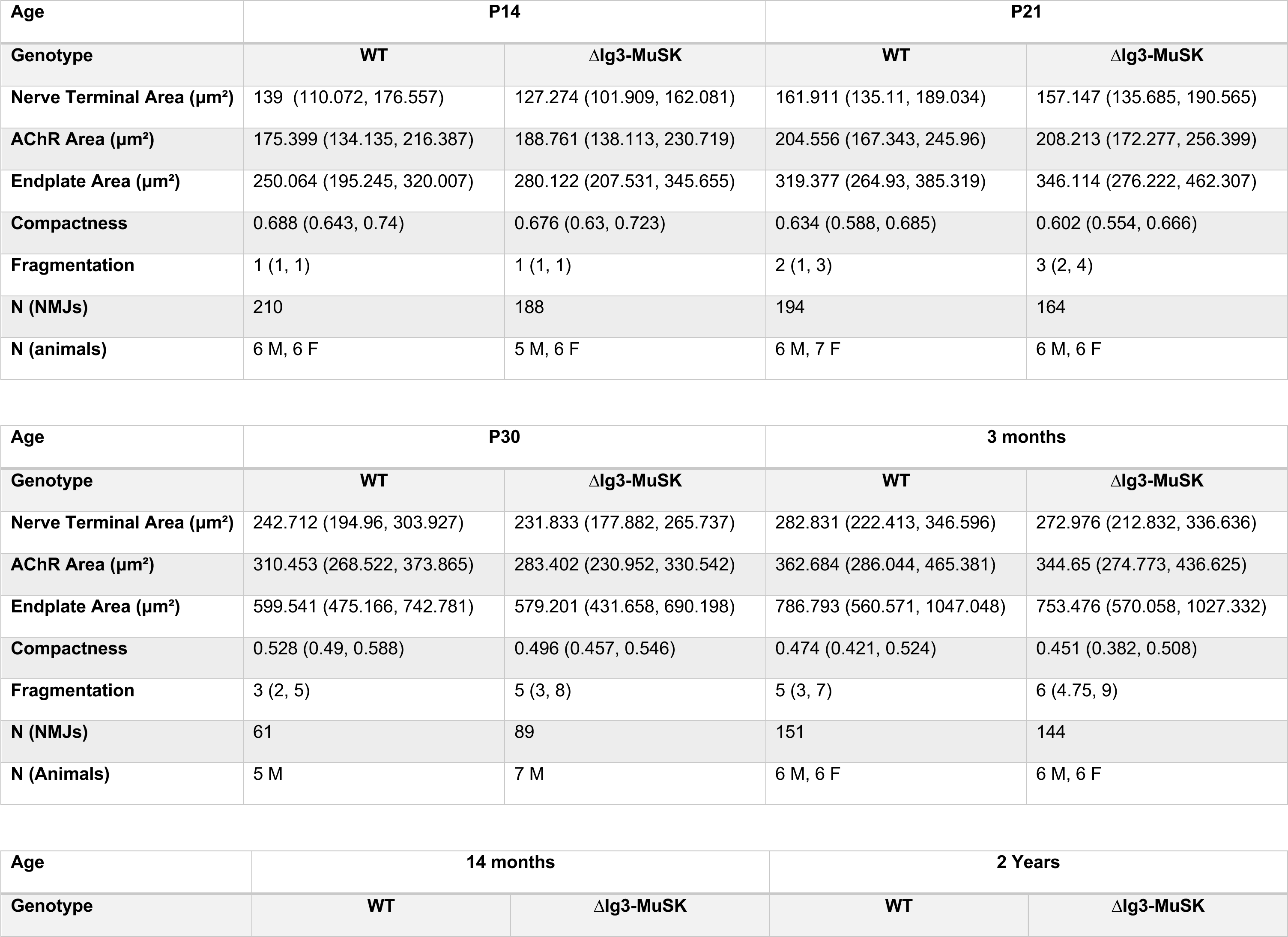
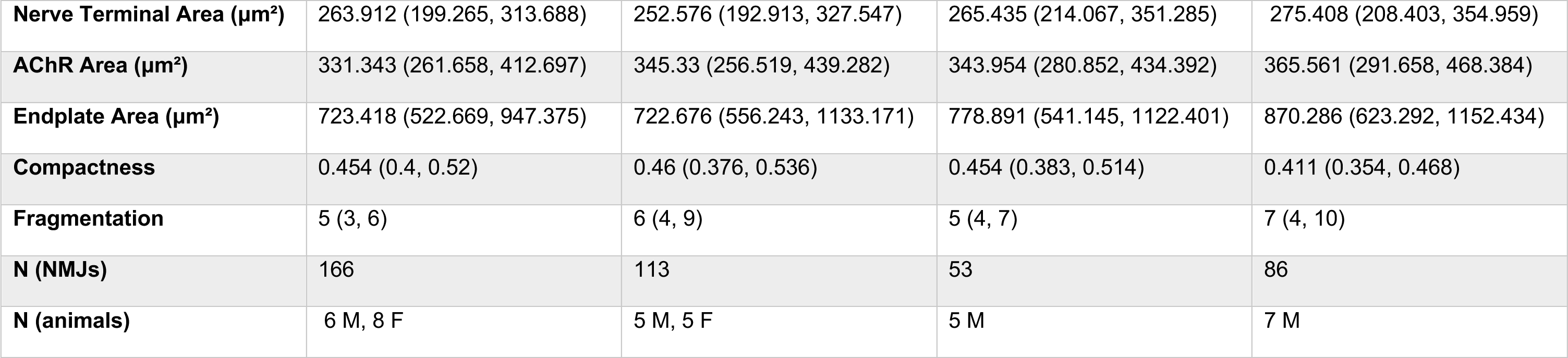
Sternomastoid NMJ morphometry data across lifespan – mixed male and female dataset. Data for each timepoint studied presented as: Median value (interquartile range). Sample sizes (animals and synapses) indicated for each timepoint. Note males only were used at P30 and 2 years; data for these ages is from the male-only dataset in Table 1-1.

| Age | P14 |  | P21 |  |
| --- | --- | --- | --- | --- |
| Genotype | WT | ΔIg3-MuSK | WT | ΔIg3-MuSK |
| Nerve Terminal Area ($\mu\text{m}^2$) | 139 (110.072, 176.557) | 127.274 (101.909, 162.081) | 161.911 (135.11, 189.034) | 157.147 (135.685, 190.565) |
| AChR Area ($\mu\text{m}^2$) | 175.399 (134.135, 216.387) | 188.761 (138.113, 230.719) | 204.556 (167.343, 245.96) | 208.213 (172.277, 256.399) |
| Endplate Area ($\mu\text{m}^2$) | 250.064 (195.245, 320.007) | 280.122 (207.531, 345.655) | 319.377 (264.93, 385.319) | 346.114 (276.222, 462.307) |
| Compactness | 0.688 (0.643, 0.74) | 0.676 (0.63, 0.723) | 0.634 (0.588, 0.685) | 0.602 (0.554, 0.666) |
| Fragmentation | 1 (1, 1) | 1 (1, 1) | 2 (1, 3) | 3 (2, 4) |
| N (NMJs) | 210 | 188 | 194 | 164 |
| N (animals) | 6 M, 6 F | 5 M, 6 F | 6 M, 7 F | 6 M, 6 F |

| Age | P30 |  | 3 months |  |
| --- | --- | --- | --- | --- |
| Genotype | WT | ΔIg3-MuSK | WT | ΔIg3-MuSK |
| Nerve Terminal Area ($\mu\text{m}^2$) | 242.712 (194.96, 303.927) | 231.833 (177.882, 265.737) | 282.831 (222.413, 346.596) | 272.976 (212.832, 336.636) |
| AChR Area ($\mu\text{m}^2$) | 310.453 (268.522, 373.865) | 283.402 (230.952, 330.542) | 362.684 (286.044, 465.381) | 344.65 (274.773, 436.625) |
| Endplate Area ($\mu\text{m}^2$) | 599.541 (475.166, 742.781) | 579.201 (431.658, 690.198) | 786.793 (560.571, 1047.048) | 753.476 (570.058, 1027.332) |
| Compactness | 0.528 (0.49, 0.588) | 0.496 (0.457, 0.546) | 0.474 (0.421, 0.524) | 0.451 (0.382, 0.508) |
| Fragmentation | 3 (2, 5) | 5 (3, 8) | 5 (3, 7) | 6 (4.75, 9) |
| N (NMJs) | 61 | 89 | 151 | 144 |
| N (Animals) | 5 M | 7 M | 6 M, 6 F | 6 M, 6 F |

**Table 2-1. Sternomastoid NMJ morphometry data across lifespan – mixed male and female dataset.**
| Age | 14 months |  | 2 Years |  |
| --- | --- | --- | --- | --- |
| Genotype | WT | ΔIg3-MuSK | WT | ΔIg3-MuSK |
| **Nerve Terminal Area ($\mu\text{m}^2$)** | 263.912 (199.265, 313.688) | 252.576 (192.913, 327.547) | 265.435 (214.067, 351.285) | 275.408 (208.403, 354.959) |
| **AChR Area ($\mu\text{m}^2$)** | 331.343 (261.658, 412.697) | 345.33 (256.519, 439.282) | 343.954 (280.852, 434.392) | 365.561 (291.658, 468.384) |
| **Endplate Area ($\mu\text{m}^2$)** | 723.418 (522.669, 947.375) | 722.676 (556.243, 1133.171) | 778.891 (541.145, 1122.401) | 870.286 (623.292, 1152.434) |
| **Compactness** | 0.454 (0.4, 0.52) | 0.46 (0.376, 0.536) | 0.454 (0.383, 0.514) | 0.411 (0.354, 0.468) |
| **Fragmentation** | 5 (3, 6) | 6 (4, 9) | 5 (4, 7) | 7 (4, 10) |
| **N (NMJs)** | 166 | 113 | 53 | 86 |
| **N (animals)** | 6 M, 8 F | 5 M, 5 F | 5 M | 7 M |

**Table 2-2.** Complete sternomastoid statistical results across lifespan-mixed male and female dataset. Best fit distribution used to fit GLM for each morphological measurement and associated p-values for genotype, sex, and interaction effects as applicable. GLMs fit by minimization of AIC. Significant p-values in bold. Note males only were used at P30 and 2 years; data for these ages is from the male-only dataset in Table 1-2.

| Measurement | Statistics | P14 | P21 | P30 | 3 months | 14 months | 2 years |
| --- | --- | --- | --- | --- | --- | --- | --- |
| **Nerve Terminal Area** | Distribution | gamma | gamma | gamma | gamma | gamma | gamma |
|  | Genotype P Value | 0.699 | 0.0655 | 0.0652 | **0.0391** | 0.1819 | 0.867 |
|  | Sex P Value | 0.508 | 0.3191 | N/A | **9.43e-07** | **0.0046** | N/A |
|  | Interaction P Value | 0.110 | **0.0173** | N/A | **0.0564** | **0.0085** | N/A |
| **AChR Area** | Distribution | gamma | gamma | gamma | gamma | gamma | gamma |
|  | Genotype P Value | **0.0497** | **0.0184** | **0.0164** | **0.000927** | 0.2059 | 0.344 |
|  | Sex P Value | 0.9140 | 0.1575 | N/A | **7.23e-10** | **5.94e-05** | N/A |
|  | Interaction P Value | 0.0960 | 0.2809 | N/A | **0.001046** | **0.0063** | N/A |
| **Endplate Area** | Distribution | gamma | gamma | gamma | gamma | gamma | gamma |
|  | Genotype P Value | 0.107 | **0.00194** | 0.637 | 0.05626 | 0.4426 | 0.281 |
|  | Sex P Value | 0.820 | 0.47118 | N/A | **5.3e-12** | **3.29e-05** | N/A |
|  | Interaction P Value | 0.721 | 0.50187 | N/A | **0.00232** | **0.0177** | N/A |
| **Compactness** | Distribution | beta | beta | beta | beta | beta | beta |
|  | Genotype P Value | **0.0129** | **0.0066** | **0.00192** | 0.334 | 0.545 | **0.0135** |
|  | Sex P Value | **0.0432** | 0.1530 | N/A | **1.44e-09** | 0.144 | N/A |
|  | Interaction P Value | **4.82e-06** | 0.9476 | N/A | 0.310 | 0.562 | N/A |
| **Fragmentation** | Distribution | poisson | poisson | negative binomial | negative binomial | negative binomial | negative binomial |
|  | Genotype P Value | 0.550 | **6.18e-05** | **1.89e-05** | **0.032187** | **5.87e-05** | **0.0236** |
|  | Sex P Value | 0.641 | 0.949 | N/A | **0.000114** | 0.909 | N/A |
|  | Interaction P Value | 0.899 | 0.843 | N/A | **0.016528** | 0.413 | N/A |

**Table 3-1.** Soleus and EDL morphometric measurements. Data for each muscle presented as: Median value (interquartile range). Sample sizes (animals and synapses) indicated for each muscle.

| **Muscle** | **Soleus** |  | **EDL** |  |
| --- | --- | --- | --- | --- |
| **Genotype** | WT | ΔIg3-MuSK | WT | ΔIg3-MuSK |
| **Nerve Terminal Area ($\mu\text{m}^2$)** | 286.639 (234.333, 383.998) | 266.081 (197.828, 331.955) | 206.099 (156.057, 231.682) | 204.254 (161.92, 295.371) |
| **AChR Area ($\mu\text{m}^2$)** | 354.355 (292.394, 455.404) | 315.762 (258.259, 387.16) | 285.985 (216.064, 317.574) | 269.352 (212.474, 358.222) |
| **Endplate Area ($\mu\text{m}^2$)** | 603.475 (497.259, 794.594) | 585.131 (466.019, 713.302) | 496.429 (370.812, 614.472) | 580.526 (387.244, 690.277) |
| **Compactness (AChR/Endplate Area)** | 0.581 (0.52, 0.637) | 0.567 (0.512, 0.628) | 0.576 (0.541, 0.641) | 0.509 (0.464, 0.586) |
| **Fragmentation** | 2 (1, 3) | 2 (1, 3) | 2 (1, 2) | 2 (2, 3) |
| **Nerve terminal caliber ($\mu\text{m}$)** | 3.217 (2.867, 3.446) | 3.397 (2.986, 3.667) | 2.31 (2.043, 2.578) | 2.449 (2.152, 2.602) |
| **Sample Size (NMJs)** | 69 | 71 | 41 | 39 |
| **Sample Size (mice)** | 6 | 6 | 3 | 3 |

**Table 3-2.** Soleus and EDL statistics. Best fit distribution used to fit generalized linear model for each morphological measurement and associated p-value for genotype. GLMs fit by minimization of AIC. Significant p-values in bold. Note: STM 3-month male nerve terminal caliber measurements can be seen in Table 1-1. Statistical results: p=0.828, GLM: gamma distribution.

| Measurement | Statistic | Soleus | EDL |
| --- | --- | --- | --- |
| Nerve Terminal area | Distribution | gamma | gamma |
|  | Genotype P Value | **0.0286** | 0.0617 |
| AChR Area | Distribution | gamma | gamma |
|  | Genotype P Value | **0.0174** | 0.376 |
| Endplate Area | Distribution | gamma | gamma |
|  | Genotype P Value | 0.113 | **0.0299** |
| Compactness | Distribution | beta | beta |
|  | Genotype P Value | 0.386 | **5.44e-5** |
| Fragmentation | Distribution | Poisson | Poisson |
|  | Genotype P Value | 0.0525 | **0.00369** |
| Nerve terminal caliber | Distribution | gamma | lognormal |
|  | Genotype P Value | **0.00575** | 0.312 |

## References

Arnold WD, Sheth KA, Wier CG, Kissel JT, Burghes AH, Kolb SJ (2015) Electrophysiological Motor Unit Number Estimation (MUNE) Measuring Compound Muscle Action Potential (CMAP) in Mouse Hindlimb Muscles. J Vis Exp Jove:52899.

Balci K, Turgut N, Nurlu G (2005) Normal values for single fiber EMG parameters of frontalis muscle in healthy subjects older than 70 years. Clin Neurophysiol 116:1555–1557.

Balice-Gordon R, Lichtman J (1990) In vivo visualization of the growth of pre-and postsynaptic elements of neuromuscular junctions in the mouse. J Neurosci 10:894–908.

Balice-Gordon RJ (1997) Age-related changes in neuromuscular innervation. Muscle Nerve 20:83–87.

Bányai L, Sonderegger P, Patthy L (2010) Agrin Binds BMP2, BMP4 and TGFβ1. Plos One 5:e10758.

Belhasan DC, Akaaboune M (2020) The role of the Dystrophin Glycoprotein Complex on the Neuromuscular System. Neurosci Lett 722:134833.

Bromberg MB, Scott DM, Group TAHC of the ASFSI (1994) Single fiber EMG reference values: Reformatted in tabular form. Muscle Nerve 17:820–821.

Bruneau EG, Akaaboune M (2006) The dynamics of recycled acetylcholine receptors at the neuromuscular junction in vivo. Development 133:4485–4493.

Burden SJ, Yumoto N, Zhang W (2013) The Role of MuSK in Synapse Formation and Neuromuscular Disease. Csh Perspect Biol 5:a009167.

Chugh D, Iyer CC, Wang X, Bobbili P, Rich MM, Arnold WD (2020) Neuromuscular junction transmission failure is a late phenotype in aging mice. Neurobiol Aging 86:182–190.

Cribari-Neto F, Zeileis A (2010) Beta Regression in R. J Stat Softw 34.

Fatt P, Katz B (1952) The electric activity of the motor end-plate. Proc R Soc Lond Ser B - Biol Sci 140:183–186.

Feng G, Mellor RH, Bernstein M, Keller-Peck C, Nguyen QT, Wallace M, Nerbonne JM, Lichtman JW, Sanes JR (2000) Imaging Neuronal Subsets in Transgenic Mice Expressing Multiple Spectral Variants of GFP. Neuron 28:41–51.

Feng Z, Ko C (2008) The Role of Glial Cells in the Formation and Maintenance of the Neuromuscular Junction. Ann Ny Acad Sci 1132:19–28.

Fish LA, Fallon JR (2020) Multiple MuSK signaling pathways and the aging neuromuscular junction. Neurosci Lett 731:135014.

Fuertes-Alvarez S, Izeta A (2021) Terminal Schwann Cell Aging: Implications for Age-Associated Neuromuscular Dysfunction. Aging Dis 12:494–514.

Furuta N, Ishizawa K, Shibata M, Tsukagoshi S, Nagamine S, Makioka K, Fujita Y, Ikeda M, Yoshimura S, Motomura M, Okamoto K, Ikeda Y (2015) Anti-MuSK Antibody-positive Myasthenia Gravis Mimicking Amyotrophic Lateral Sclerosis. Internal Med 54:2497–2501.

Gilmore KJ, Morat T, Doherty TJ, Rice CL (2017) Motor unit number estimation and neuromuscular fidelity in 3 stages of sarcopenia. Muscle Nerve 55:676–684.

Glass DJ, Bowen DC, Stitt TN, Radziejewski C, Bruno J, Ryan TE, Gies DR, Shah S, Mattsson K, Burden SJ, DiStefano PS, Valenzuela DM, DeChiara TM, Yancopoulos GD (1996) Agrin Acts via a MuSK Receptor Complex. Cell 85:513–523.

Haddix SG, Lee Y il, Kornegay JN, Thompson WJ (2018) Cycles of myofiber degeneration and regeneration lead to remodeling of the neuromuscular junction in two mammalian models of Duchenne muscular dystrophy. PLoS ONE 13:e0205926.

Huijbers MG, Niks EH, Klooster R, Visser M de, Kuks JB, Veldink JH, Klarenbeek P, Damme PV, Baets MH de, Maarel SM van der, Berg LH van den, Verschuuren JJ (2016) Myasthenia gravis with muscle specific kinase antibodies mimicking amyotrophic lateral sclerosis. Neuromuscular Disord 26:350–353.

Huijbers MG, Zhang W, Klooster R, Niks EH, Friese MB, Straasheijm KR, Thijssen PE, Vrolijk H, Plomp JJ, Vogels P, Losen M, Maarel SMV der, Burden SJ, Verschuuren JJ (2013) MuSK IgG4 autoantibodies cause myasthenia gravis by inhibiting binding between MuSK and Lrp4. Proc National Acad Sci 110:20783–20788.

Jaime D, Fish LA, Madigan LA, Ewing ME, Fallon JR (2022) The MuSK-BMP pathway maintains myofiber size in slow muscle through regulation of Akt-mTOR signaling.

Jaime D, Fish LA, Madigan LA, Xi C, Piccoli G, Ewing MD, Blaauw B, Fallon JR (2023) The MuSK-BMP pathway maintains myofiber size in slow muscle through regulation of Akt-mTOR signaling. Skeletal Muscle (in press).

Jones RA, Reich CD, Dissanayake KN, Kristmundsdottir F, Findlater GS, Ribchester RR, Simmen MW, Gillingwater TH (2016) NMJ-morph reveals principal components of synaptic morphology influencing structure–function relationships at the neuromuscular junction. Open Biol 6:160240.

Kang H, Tian L, Mikesh M, Lichtman JW, Thompson WJ (2014) Terminal Schwann Cells Participate in Neuromuscular Synapse Remodeling during Reinnervation following Nerve Injury. J Neurosci 34:6323–6333.

Lee Y il, Thompson WJ, Harlow ML (2017) Schwann cells participate in synapse elimination at the developing neuromuscular junction. Curr Opin Neurobiol 47:176–181.

Li Y, Lee Y i, Thompson WJ (2011) Changes in Aging Mouse Neuromuscular Junctions Are Explained by Degeneration and Regeneration of Muscle Fiber Segments at the Synapse. J Neurosci 31:14910–14919.

Lichtman J, Magrassi L, Purves D (1987) Visualization of neuromuscular junctions over periods of several months in living mice. J Neurosci 7:1215–1222.

Lømo T, Waerhaug O (1985) Motor endplates in fast and slow muscles of the rat: what determines their difference? J Physiol 80:290–297.

Madigan LA, Jaime D, Fallon JR (2023) MuSK-BMP signaling in adult muscle stem cells maintains quiescence and regulates myofiber size. bioRxiv:2023.05.17.541238.

Minty G, Hoppen A, Boehm I, Alhindi A, Gibb L, Potter E, Wagner BC, Miller J, Skipworth RJE, Gillingwater TH, Jones RA (2020) aNMJ-morph: a simple macro for rapid analysis of neuromuscular junction morphology. Roy Soc Open Sci 7:200128.

Mitchell WK, Williams J, Atherton P, Larvin M, Lund J, Narici M (2012) Sarcopenia, Dynapenia, and the Impact of Advancing Age on Human Skeletal Muscle Size and Strength; a Quantitative Review. Front Physiol 3:260.

Padilla CJ, Harrigan ME, Harris H, Schwab JM, Rutkove SB, Rich MM, Clark BC, Arnold WD (2021) Profiling age-related muscle weakness and wasting: neuromuscular junction transmission as a driver of age-related physical decline. Geroscience 43:1265–1281.

Poort JE, Rheuben MB, Breedlove SM, Jordan CL (2016) Neuromuscular junctions are pathological but not denervated in two mouse models of spinal bulbar muscular atrophy. Hum Mol Genet 25:3768–3783.

Rudolf R, Khan MM, Labeit S, Deschenes MR (2014) Degeneration of Neuromuscular Junction in Age and Dystrophy. Front Aging Neurosci 6:99.

Sanes JR, Lichtman JW (1999) DEVELOPMENT OF THE VERTEBRATE NEUROMUSCULAR JUNCTION. Annu Rev Neurosci 22:389–442.

Sanes JR, Lichtman JW (2001) Induction, assembly, maturation and maintenance of a postsynaptic apparatus. Nat Rev Neurosci 2:791–805.

Schiaffino S, Reggiani C (2011) Fiber Types in Mammalian Skeletal Muscles. Physiol Rev 91:1447–1531.

Sharp AA, Caldwell JH (1996) Aggregation of Sodium Channels Induced by a Postnatally Upregulated Isoform of Agrin. J Neurosci 16:6775–6783.

Sheth KA, Iyer CC, Wier CG, Crum AE, Bratasz A, Kolb SJ, Clark BC, Burghes AHM, Arnold WD (2018) Muscle strength and size are associated with motor unit connectivity in aged mice. Neurobiol Aging 67:128–136.

Shi L, Fu AKY, Ip NY (2012) Molecular mechanisms underlying maturation and maintenance of the vertebrate neuromuscular junction. Trends Neurosci 35:441–453.

Slater CR (2008) Structural Factors Influencing the Efficacy of Neuromuscular Transmission. Ann Ny Acad Sci 1132:1–12.

Slater CR (2017) The Structure of Human Neuromuscular Junctions: Some Unanswered Molecular Questions. Int J Mol Sci 18:2183.

Slater CR (2020) ‘Fragmentation’ of NMJs: a sign of degeneration or regeneration? A long journey with many junctions. Neuroscience 439:28–40.

Stiegler AL, Burden SJ, Hubbard SR (2006) Crystal Structure of the Agrin-responsive Immunoglobulin-like Domains 1 and 2 of the Receptor Tyrosine Kinase MuSK. J Mol Biol 364:424–433.

Team RC (2021) R: A language and environment for statistical computing. Vienna, Austria: R Foundation for Statistical Computing. Available at: https://www.R-project.org/.

Tintignac LA, Brenner H-R, Rüegg MA (2015) Mechanisms Regulating Neuromuscular Junction Development and Function and Causes of Muscle Wasting. Physiol Rev 95:809–852.

Valdez G, Tapia JC, Kang H, Clemenson GD, Gage FH, Lichtman JW, Sanes JR (2010) Attenuation of age-related changes in mouse neuromuscular synapses by caloric restriction and exercise. Proc National Acad Sci 107:14863–14868.

Valdez G, Tapia JC, Lichtman JW, Fox MA, Sanes JR (2012) Shared Resistance to Aging and ALS in Neuromuscular Junctions of Specific Muscles. Plos One 7:e34640.

Venables W, Ripley B (2002) Modern Applied Statistics with S, Forth. New York: Springer. Available at: https://www.stats.ox.ac.uk/pub/MASS4/.

Wang X, Engisch KL, Li Y, Pinter MJ, Cope TC, Rich MM (2004) Decreased Synaptic Activity Shifts the Calcium Dependence of Release at the Mammalian Neuromuscular Junction In Vivo. J Neurosci 24:10687–10692.

Watty A, Neubauer G, Dreger M, Zimmer M, Wilm M, Burden SJ (2000) The in vitro and in vivo phosphotyrosine map of activated MuSK. Proc National Acad Sci 97:4585–4590.

Wickham H (2016) ggplot2, Elegant Graphics for Data Analysis.:109–145.

Wickham H et al. (2019) Welcome to the Tidyverse. J Open Source Softw 4:1686.

Wilke CO (2020) cowplot: Streamlined Plot Theme and Plot Annotations for “ggplot2.” Available at: https://CRAN.R-project.org/package=cowplot.

Willadt S, Nash M, Slater CR (2016) Age-related fragmentation of the motor endplate is not associated with impaired neuromuscular transmission in the mouse diaphragm. Sci Rep 6:24849.

Wood SJ, Slater CR (2001) Safety factor at the neuromuscular junction. Prog Neurobiol 64:393–429.

Yilmaz A, Kattamuri C, Ozdeslik RN, Schmiedel C, Mentzer S, Schorl C, Oancea E, Thompson TB, Fallon JR (2016) MuSK is a BMP co-receptor that shapes BMP responses and calcium signaling in muscle cells. Sci Signal 9:ra87.

Yumoto N, Kim N, Burden SJ (2012) Lrp4 Is A Retrograde Signal For Presynaptic Differentiation At Neuromuscular Synapses. Nature 489:438–442.

Zhang C, Joshi A, Liu Y, Sert O, Haddix SG, Teliska LH, Rasband A, Rodney GG, Rasband MN (2021) Ankyrin-dependent Na+ channel clustering prevents neuromuscular synapse fatigue. Curr Biol 31:3810–3819.e4.

Zhang W, Coldefy A-S, Hubbard SR, Burden SJ (2011) Agrin Binds to the N-terminal Region of Lrp4 Protein and Stimulates Association between Lrp4 and the First Immunoglobulin-like Domain in Muscle-specific Kinase (MuSK)*. J Biol Chem 286:40624–40630.

Zong Y, Zhang B, Gu S, Lee K, Zhou J, Yao G, Figueiredo D, Perry K, Mei L, Jin R (2012) Structural basis of agrin–LRP4–MuSK signaling. Gene Dev 26:247–258.

